# Network-mediated diffusion produces disordered self-organization in vegetation^⋆^

**DOI:** 10.64898/2026.04.16.718764

**Authors:** Sara Filippini, Luca Ridolfi, Jost von Hardenberg

**Affiliations:** Politecnico di Torino, Corso Duca degli Abruzzi 24, Torino (TO), 10129, Italy; National Institute of Oceanography and Applied Geophysics, Borgo Grotta Gigante 42/C, Sgonico (TS), 34010, Italy

**Keywords:** vegetation, patterns, dryland, network, self-organization, non-local, regularity

## Abstract

Vegetation patterns in arid and semiarid regions emerge as a result of a self-organization process triggered by water scarcity. While highly regular patterns are well reproduced through reaction-diffusion models, the origin of commonly observed disordered patterns remains debated. Several studies have attributed them either to spatial heterogeneity, or to a global competition regime induced by domain scale water transport. Here, we propose a new model in which biomass and water diffuse through networks constructed by adding random links (shortcuts) onto a regular lattice. By reproducing the naturally occurring spatially heterogeneous co-existence of different transport scales, our model incorporates both aspects of previous explanations for the origin of disordered patterns. It also allows us to analyze our results through the lens of network theory. In fact, as the density of shortcuts increases, the diffusion networks transition from a regular lattice through a small world network to a random network, resulting in different pattern formation behaviors. On a regular lattice, high-regularity patterns develop reflecting local diffusion processes. On a random network, the system is dominated by domain scale diffusion yielding either a uniform state or a single patch. In the intermediate shortcut density range, on a small world network topology, the interaction between the two scales of diffusion generates two kinds of disordered patterns, both coherent with observations: low-regularity patterns with a well-defined characteristic wavelength, and irregular patterns characterized by a broad patch size distribution. Our model predicts that these two pattern types would vary significantly in their resilience to environmental pressure.

**Highlights:**

- In real ecosystems, different water and biomass transport scales often co-exist
- We use network diffusion to simulate long-distance transport in a vegetation model
- Disordered patterns emerge when long-distance transport is sparingly available
- Low-regularity patterns when long-distance transport of biomass is possible
- Irregular broad patch size patterns when long-distance transport of water is possible

## 1. Introduction

In many biological systems subject to resource limitations, spatial patterns emerge as the result of a self-organization process (Halatek et al., 2018; Meron, 2015; Rietkerk and van De Koppel, 2008). Several classes of dynamical systems, in fact, exhibit regions of the parameter space where the uniform solution becomes unstable, and the system naturally evolves into stationary, non-uniform patterns (Cross and Hohenberg, 1993). This is the case of activator-inhibitor systems subject to the well-known Turing instability. Traditionally applied to reaction-diffusion chemical systems, the Turing pattern formation mechanism has been extended to a number of other contexts (Koch and Meinhardt, 1994; Maini et al., 1997; Turing, 1952).

One of its most noticeable applications concerns dryland vegetation. Regular spatial patterns are commonly observed in the vegetation of arid and semi-arid regions, often unaccounted for by topography or local soil properties (D’Odorico et al., 2006; von Hardenberg et al., 2001; Meron, 2015). They may be explained as a result of the balance between short-range facilitation and long-range competition between plants, producing a Turing-like instability (Borgogno et al., 2009; Martinez-Garcia et al., 2023; Meron, 2018; Rietkerk et al., 2002). In this case, biomass acts as the activator and lack of water (dryness) as the inhibitor. Biomass promotes further biomass growth locally through mutualistic processes such as evaporation reduction through shading, increased water uptake due to root expansion, and increased soil infiltration through the prevention of soil crust formation. As water is consumed by growing plants, biomass also promotes local increases in dryness. In a water limited ecosystem, dryness limits biomass growth. Since water travels significantly faster than biomass, mutualistic processes act on smaller spatial scales than competition for the limited water resource, leading to a Turing-like instability below a certain precipitation threshold (D’Odorico et al., 2006; von Hardenberg et al., 2010; Rietkerk et al., 2002). This instability often increases ecosystem resilience by enabling the system to avoid a catastrophic transition from uniform vegetation to desert (Rietkerk et al., 2021). By simulating the processes we outlined, several vegetation models have been able to reproduce highly regular patterns (Borgogno et al., 2009).

However, real-world patterns commonly display a significantly lower regularity than model predictions (Kästner et al., 2024b; Pinto-Ramos et al., 2023; Pinto-Ramos et al., 2025), and entirely irregular patterns characterized by wide patch size distributions have been observed in various environments (Kéfi et al., 2007, 2011; Lin et al., 2010; Maestre and Escudero, 2009; Scanlon et al., 2007). Some recent studies explain this lack of regularity as the result of environmental heterogeneity, and reproduce disordered patterns through the inclusion of spatially varying parameters in the vegetation models (Kästner et al., 2025; Pinto-Ramos et al., 2022; Pinto-Ramos et al., 2025; Yizhaq et al., 2014). Yet these models are rarely capable of reproducing the wide patch size distributions observed in the field (Kästner et al., 2025; Pinto-Ramos et al., 2025; Yizhaq et al., 2014). Pinto-Ramos et al. (2022) find power-law clustering in a model with spatially varying mortality rate, but only as a phase separation pattern, independent of the Turing instability. Furthermore, this approach is limited in its generality due to its dependence on the choice of the varying parameter. At the same time, irregular, wide patch size distribution patterns emerge as transient states in spatially homogeneous systems in which the fast spatial distribution of water leads to a global competition regime (von Hardenberg et al., 2010; Kletter et al., 2012). Thus, two unrelated mechanisms are proposed as explanation of the same phenomenon, each with some limitations in their results.

We propose here a different approach incorporating both of these aspects (global scale diffusion and spatial heterogeneity), inspired by recent work in the field of pattern formation on networks (Imayama and Shiwa, 2008; McCullen and Wagenknecht, 2016; Nakao and Mikhailov, 2010). We increase transport of either water or biomass through the system by gradually increasing the connectivity of the relative transport networks, rather than modifying the diffusion constants, thus accounting for transport mechanisms acting over different spatial scales in a spatially heterogeneous manner (see section 2.2). We then study the evolution of the pattern formation process over different network topologies, revealing the emergence of different kinds of stable low-regularity and irregular patterns in an intermediate connectivity range. Our approach allows us to apply all the tools developed within the field of network theory in order to interpret our model results.

## 2. Model

### 2.1. Differential equations statement

We adopted the model formulated by Zelnik et al. (2015), which is a reduced version the one by Gilad et al. (Gilad et al., 2004; von Hardenberg et al., 2010). The model consists of two coupled first-order differential equations, that describe the dynamics of water and biomass densities in a 2-dimensional space. Biomass grows as a logistic curve approaching the carrying capacity of each spatial point, the growth rate being proportional to the available water, and it is subject to a linear mortality rate. Water is supplied through precipitation (the only climate variable we consider), and consumed through a linear evaporation rate and a linear vegetation uptake rate, the latter one proportional to local biomass. Both variables are subject to a linear diffusion term (assuming flat terrain, we do not include advection terms).

Two mutualistic mechanisms are explicitly included: the water evaporation rate is reduced proportionally to local biomass to simulate shading, and the biomass growth term, as well as the water consumption term, include a quadratic factor approximating the increased water uptake due to root extension (Zelnik et al., 2015).

The model equations, in dimensionless form, are as follows:

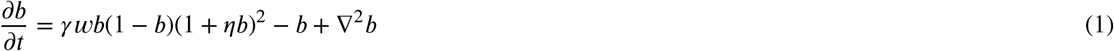

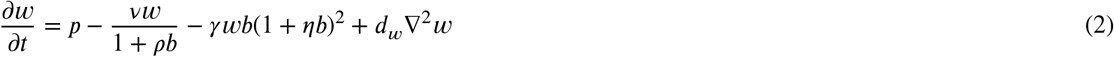

where *b* and *w* refer to biomass and water non-dimensional densities, respectively. *γ* refers to the water uptake rate by growing plants, *ν* represents the water evaporation rate, and *d*_*w*_ is the ratio of water and biomass dimensional diffusivities. Finally, *ρ* controls the intensity of evaporation reduction due to shading, and *η* the non-linear increase in water uptake due to root extension.

This model was derived from the more complex version described in Gilad et al. (Gilad et al., 2004; von Hardenberg et al., 2010), under the assumptions that the lateral root size of each plant is much smaller than the scale of pattern formation and the infiltration contrast between vegetated and bare soil areas is negligible. We chose this reduced version, in which the diffusive terms are the only non-local terms, since it simplifies modeling over a network structure and allows us to expand the scope of our results to other pattern-forming systems.

The dimensional version of equations (1-2), as well as the definition of the non-dimensional terms, can be found in the Supplementary Material (S1). The values of the parameters were adopted from Zelnik et al. (2015), and they are typical for grassy dryland vegetation. Our results are qualitatively robust to variations of the parameters choice. From here on, we will be referring only to non-dimensional variables and parameters (which we write in small caps), unless otherwise specified.

It is useful to recall the linear stability analysis of model (1-2) in reference to precipitation (*p*), in order to identify the spatially uniform solutions and their stability intervals. The dynamical system exhibits three spatially uniform steady-states, shown in figure 1 in terms of their biomass density, and in the inset on the right side of the figure in terms of their water density (water and biomass density are reported as spatially average quantities ⟨.⟩ in order to include non-uniform states). The desert solution (*b*(*x, y*) = 0 ∀ *x, y*) exists for all precipitation values, and two uniform vegetation solutions (*b*(*x, y*) = *C*_1_(*p*) and *b*(*x, y*) = *C*_2_(*p*), *C*_1_(*p*) < *C*_2_(*p*)) exist for *p* ≥ 1.52. With our parameter choice, the desert state is stable for *p* < 3.12. Above this precipitation value it becomes unstable to all (uniform or non-uniform) perturbations. The uniform solution *b*(*x, y*) = *C*_1_(*p*) is always unstable to all perturbations, and we will disregard it. The uniform solution *b*(*x, y*) = *C*_2_(*p*) is stable at *p* > 1.71, and unstable to non-uniform perturbations (marginally stable) in the interval 1.52 ≤ *p* ≤ 1.71.

**Figure 1:**
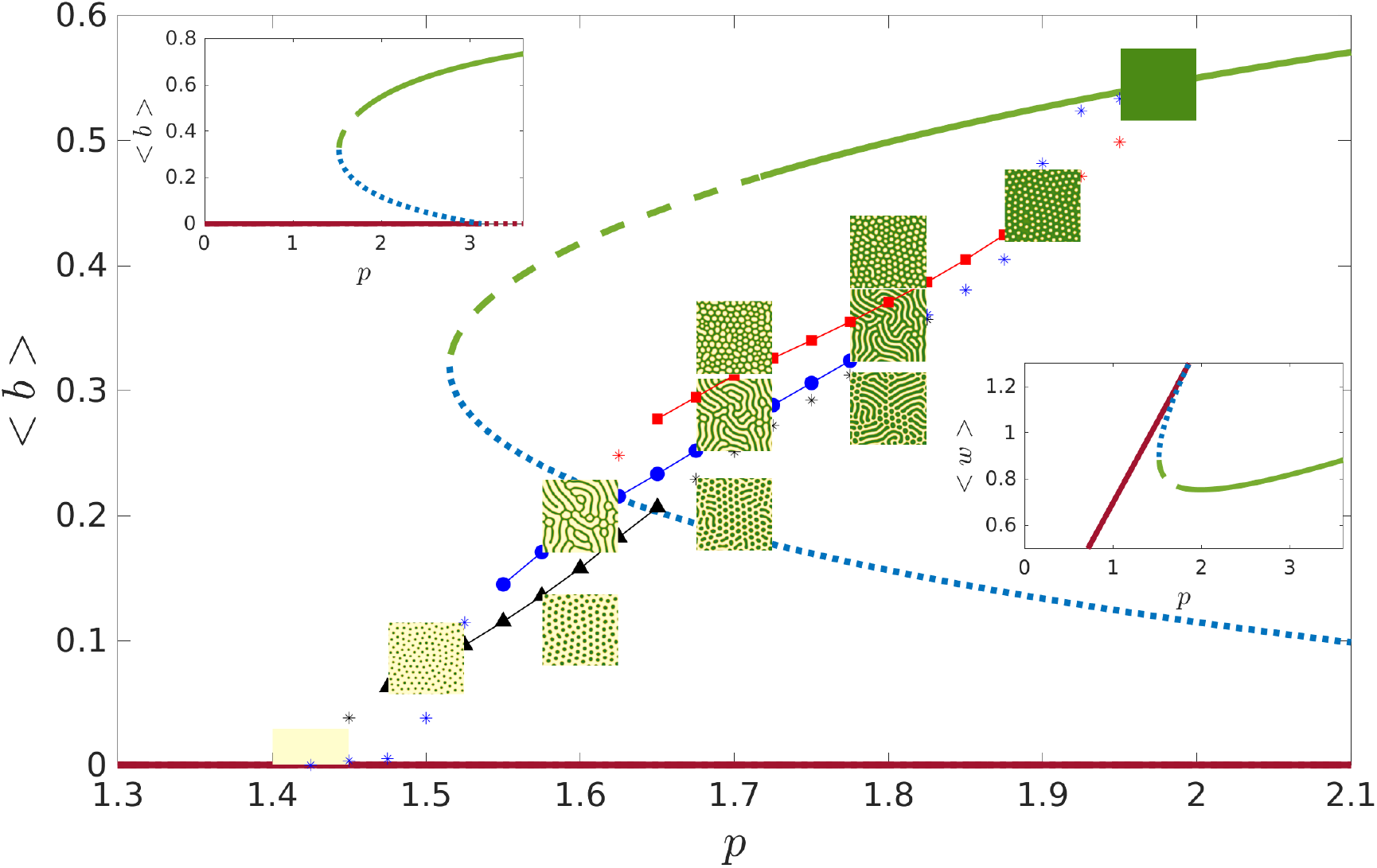
Main figure: Stability diagram of the model (1-2). The green line stands for the uniform vegetation state *b*(*x, y*) = *C*_2_(*p*), the blue line for the uniform vegetation state *b*(*x, y*) = *C*_1_(*p*), and the red line for the desert state *b*(*x, y*) = 0. Solid lines represent stable states, dotted lines unstable states and dashed lines marginally stable states (unstable to non-uniform perturbations). The connected scatter plots show the results of three different continuation series, starting from patterned solutions obtained at *p* = 1.5 (black line, triangular markers), *p* = 1.6 (blue line, circular markers) and *p* = 1.7 (red line, square markers). The asterisks represent mixed solutions. The dimensionless domain size of the simulations was 336×336, with resolution Δ*x* = 0.84 (400×400 points). The simulation time was 5000 in dimensionless time units. Top left inset: same stability diagram of the uniform solutions over a wider precipitation range. Bottom right inset: stability diagram of the uniform solutions in terms of their water density.

Besides the uniform solutions, several stable non-uniform solutions are available to the system. At precipitation values below the stability range of the uniform vegetation solution (*p* ≤ 1.71), any non-uniform perturbation of the uniform state leads to the emergence of patterned states.

We may estimate numerically the stability intervals of different patterned states through continuation technique, meaning we gradually change the precipitation using the patterned state as initial condition (Dijkstra, 2011; Krauskopf, 2007). Figure 1 shows the results of three different continuation series, starting from “dots”, “stripes” and “gaps” solutions obtained at *p* = 1.5, *p* = 1.6 and *p* = 1.7, respectively.

We observe the typical progression from “dots” to “stripes” to “gaps” patterns for increasing precipitation (Meron, 2015). We detect large co-stability intervals – i.e. precipitation intervals in which two different solutions are stable – between successive patterns, as well as a short interval of co-stability between all three patterns. Co-stability intervals between patterned states and uniform solutions are also evident. The asterisks in figure 1 refer to “mixed-solutions”: here, the same spatial domain hosts different patterns as a result of a partial transition. The fronts separating the different patterns may be stable, leading to stable “mixed-states”, or unstable, producing long transients at the end of which one of the patterns takes over the whole domain. Stable mixed-states between dots and stripes are particularly common.

Compared to the marginally stable uniform state, the patterned states have a higher mean water density and a lower mean biomass density. Thus we may interpret pattern formation as a strategy adopted by the ecosystem to form “water reserves” (the bare soil areas of the patterns) which ensure stability to perturbations, while trading off some biomass.

### 2.2. Network-mediated diffusion

Equations (1-2) may be discretized as to operate over an arbitrary network topology:

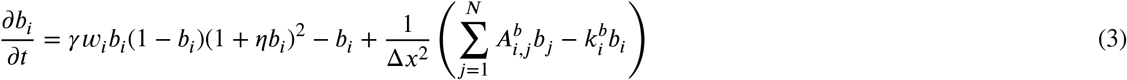

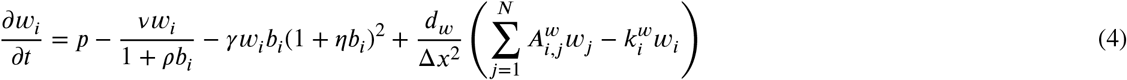

where *i* refers to a node of the network, with *N* the total number of nodes. The matrices **A**^**b**^ and **A**^**w**^ and the vectors **k**^**b**^ and **k**^**w**^ contain all the information regarding the topology of the biomass and water diffusion networks. **A** stands for the adjacency matrix of the network – i.e. *A*_*i,i*_ = 0, *A*_*i,j*_ = 1 if there is an edge between nodes *i* and *j*, otherwise *A*_*i,j*_ = 0 –, while *k*_*i*_ represents the degree of node *i* (number of neighbors). Δ*x* is the spatial resolution of the model (each edge has a weight of 1/Δ*x*^2^).

When both the biomass and the water diffusion networks are 2-dimensional lattices, equations (3-4) reduce simply to the finite difference discretizations of equations (1-2). This network topology corresponds to an assumption of homogeneous and isotropic diffusion in every point of the domain. However, in real ecosystems, overall transport of water and biomass result from the sum of different mechanisms, which may act on different spatial scales. Further, the availability of such mechanisms may be tied to local conditions, and thus be spatially heterogeneous in a non-uniform environment.

Most dryland species are capable of both vegetative (clonal) reproduction and sexual reproduction (seed/pollen dispersal) (Fukui and Araki, 2014; Hartnett et al., 2006; Saixiyala et al., 2023; Wang et al., 2018). While the spatial scale of vegetative reproduction is generally on the order of the plant size (Freschet et al., 2021), seed dispersal distances vary greatly depending on the dispersal vectors (e.g. wind, insects, birds) (Robledo-Arnuncio et al., 2014; Tamme et al., 2014; Vittoz and Engler, 2007). A common feature is a fat-tailed distribution of dispersal distances, meaning most dispersal events happen within a short range and a small number on much larger distances (Bullock and Clarke, 2000; Merwin et al., 2012; Tamme et al., 2014). The tail of the distribution is usually assessed as well above 100 meters for vector assisted dispersal (Tamme et al., 2014; Thomson et al., 2011; Vittoz and Engler, 2007). Since dispersal patterns depend on vector availability (e.g. wind patterns and trappings, insects tracks, bird nests), in non-uniform environments we may expect long-distance transport of biomass to be locally anisotropic and heterogeneous, leading to interactions between privileged points.

Similarly, in layered, spatially heterogeneous soils, different water diffusion mechanisms are active, often leading to different transport scales (Band et al., 2014; Graham and Lin, 2012; Hardie et al., 2012). Across the landscape, areas of relatively high permeability are commonly observed, for instance as the result of a localized macropore network (Buttle and McDonald, 2002; Zhang et al., 2016). In these areas, the soil characteristics allow water infiltration into deeper soil layers, as well as deeper root growth and water extraction (Behrend et al., 2025; Buttle and McDonald, 2002; Wendel et al., 2022). In these lower layers, water may exploit fast subsurface lateral redistribution mechanisms, such as diffusion along permeability contrasts or at the soil-bedrock interface, or preferential flow through soil pipes (Band et al., 2014; Buttle and McDonald, 2002; Graham and Lin, 2012; Hardie et al., 2012). Thanks to these mechanisms, subsurface flow may be orders of magnitude faster than diffusion in the upper layers through the soil matrix (Graham and Lin, 2012; Hardie et al., 2012), leading to effective interactions between distant points of the domain characterized by relatively high permeability.

In our model, we represent the co-existence of different transport scales as alterations of the diffusion networks of either biomass or water. Starting with a 2-dimensional lattice, we add randomly distributed links (shortcuts) in controlled number, while preserving all the links of the original grid, as schematized in figure 2. Or, in other words, we overlay a fully connected slow transport network (the original grid) with a sparse fast transport network (the shortcuts). From a network theory perspective, we applied a version of the Watts-Strogatz network model, which interpolates between a 2-dimensional lattice and a random network (Newman and Watts, 1999; Watts and Strogatz, 1998; Watts, 1999). The parameters *ϕ*_*b*_ and *ϕ*_*w*_ refer to the ratio between the number of shortcuts and the number of bonds in the original lattice, in the biomass and water diffusion networks, respectively. In our application of the network model, these values become a parametrization of the relative availability of fast (domain scale) vs slow (grid scale) transport.

**Figure 2:**
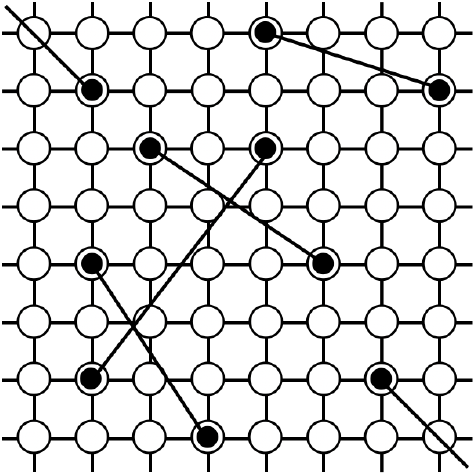
Schematic representation of the network model we employed (Newman and Watts, 1999). In this case *ϕ* = 5/128, assuming periodic boundary conditions.

A more common approach to simulating non-local dispersal is the use of a convolution integral term (Eigentler and Sherratt, 2018; Pueyo et al., 2008). Compared to this method, here we account for spatial variability in the availability of transport mechanisms, and we avoid an explicit integro-differential model, thus significantly reducing the computational burden and allowing for more extensive numerical analysis. We also gain the advantage of a direct analogy with network theory. This comes at the cost of an important simplification in the representation of transport distances, essentially reducing a fat-tailed distribution to only two scales (grid scale and domain scale).

Two main assumptions underpin the version of the model presented here. Firstly, we assume that the tail of the transport distances represented by the added shortcuts is much larger than domain size, so that the shortcuts may be randomly distributed over the domain. In our simulations, we used a square domain of ≈ 100 meters side, within the range of the mechanisms outline above. Furthermore, we neglect the time dependence of both diffusion networks. Both assumptions may be challenged and the model may be extended to relax them, but as a first step in considering this new network-based modeling method, we take a minimalist approach in terms of the (hyper)parameters.

We limit our analysis so that each point may have at most 5 neighbors (each spatial point participates in the fast transport network), hence *ϕ*_*b*_ and *ϕ*_*w*_ range between 0 and 0.25. We further restrict our study to the case of *ϕ*_*b*_*ϕ*_*w*_ = 0, studying separately the effect of biomass or water diffusion over shortcut-augmented networks.

## 3. Results

### 3.1. Biomass diffuses through a shortcut-augmented network (ϕ_b_ > 0, ϕ_w_ = 0)

In this section we compare the results obtained at different values of the parameter *ϕ*_*b*_, while keeping *ϕ*_*w*_ = 0, meaning we examine the impact on the system of an increasingly dense long-distance biomass transport network. Figure 3 shows the final stage of 21 simulations, at 3 different precipitation values and 7 different *ϕ*_*b*_ values. The initial biomass distribution for all simulations was uniform vegetation perturbed with uniformly distributed random noise in the interval [−0.05, 0.05]. We chose a dimensionless simulation time of 5000, as the patterns approached asymptotic stability around this value.

**Figure 3:**
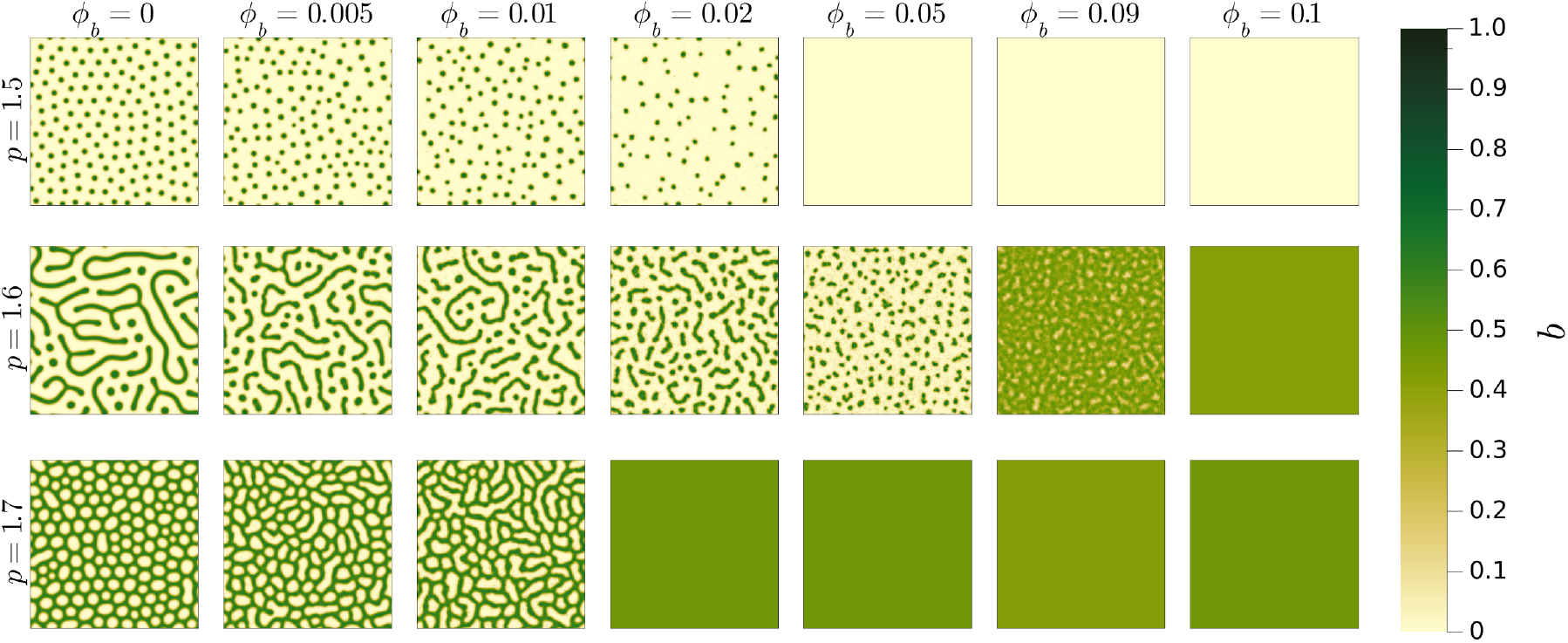
Biomass density distribution resulting from 21 different simulations at different *ϕ*_*b*_ and *p* values. The initial condition for all simulations is a uniform state perturbed with uniformly distributed random noise in the interval [−0.05, 0.05]. The simulation length is 5000 dimensionless times and the dimensionless domain size is 336×336 (400×400 points, spatial resolution 0.84).

Firstly, we observe that, at all precipitation values, the patterns give way to uniform solutions at sufficiently high shortcut densities. At *p* = 1.5, the desert state is the only stable solution for *ϕ*_*b*_ ≥ 0.05. At *p* = 1.6 and *p* = 1.7, the uniform vegetation solution becomes stable for *ϕ*_*b*_ ≥ 0.1 and *ϕ*_*b*_ ≥ 0.02, respectively. We may deduce that the addition of shortcuts alters the precipitation range in which we observe pattern formation, compared to the case of a regular lattice depicted in figure 1. To investigate this behavior, we run simulations across a range of *p* and *ϕ*_*b*_, preserving the domain size, simulation time, and initial condition as in figure 3, and map the resulting final states in figure 4. From here on we define any stable spatially heterogeneous distribution of biomass as patterns. We notice that the precipitation value below which the uniform solution becomes unstable to non-uniform perturbations (*p*_1_) decreases for increasing *ϕ*_*b*_. The lowest precipitation value at which patterns spontaneously emerge (*p*_2_) increases for increasing *ϕ*_*b*_, reaching a peak at *ϕ*_*b*_ ∼ 0.1, and later decreasing until *ϕ*_*b*_ ∼ 0.15. Thus, the precipitation range in which the system self-organizes into patterns in response to a disturbance of the uniform state reduces for increasing *ϕ*_*b*_. At high shortcut densities (*ϕ*_*b*_ ≳ 0.15), pattern formation is inhibited, and the system evolves towards either the uniform vegetation solution or the desert state.

**Figure 4:**
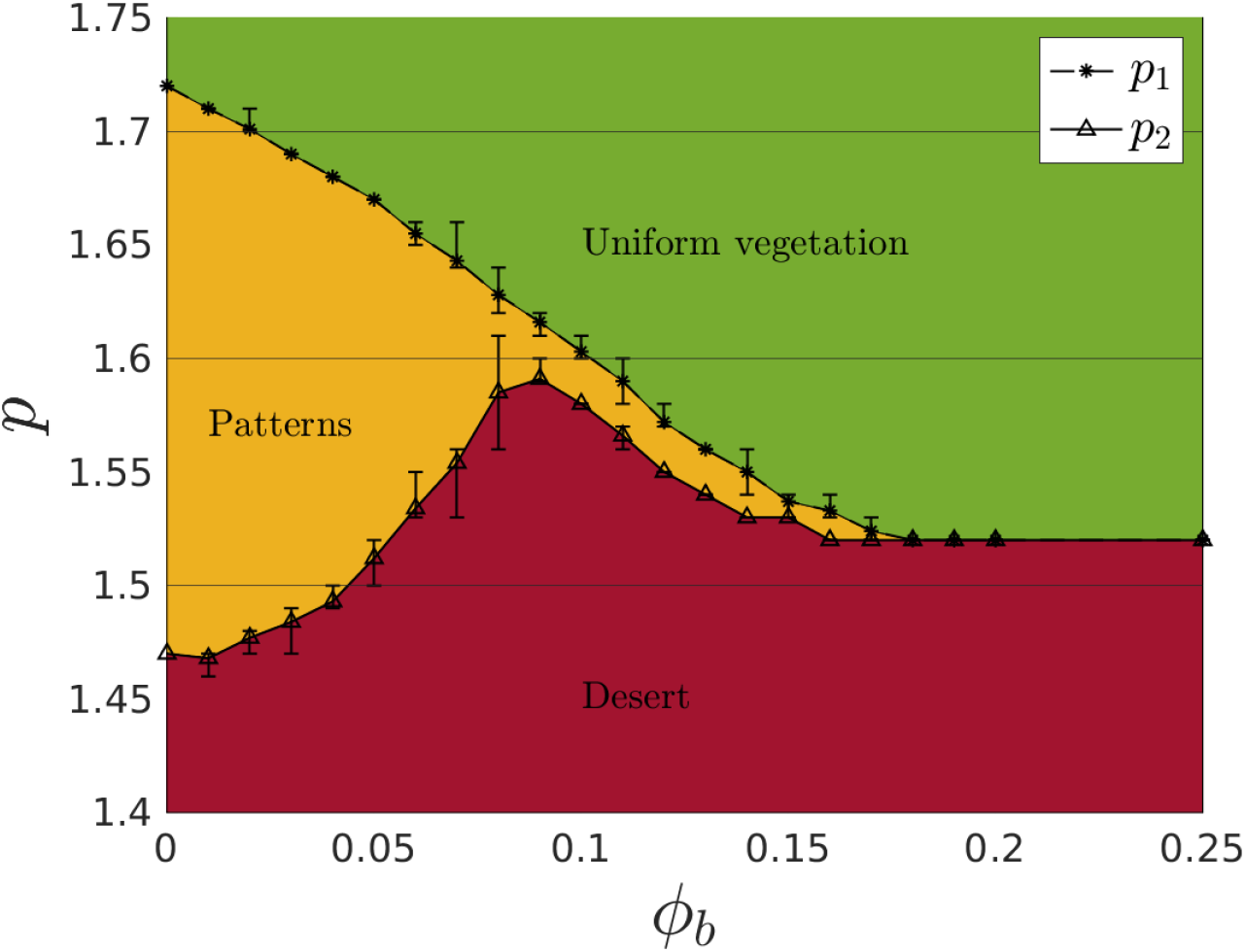
Diagram showing the final stage of simulations at different values of *p* and *ϕ*_*b*_, with initial condition uniform vegetation perturbed with uniformly distributed random noise in the interval [−0.05, 0.05]. The simulation time and domain size are the same as in figure 3. For *p* ≥ *p*_1_ the uniform vegetation state is stable, and the perturbation is suppressed. For *p* < *p*_2_ the system evolves into a desert state. In the interval *p*_1_ > *p* ≥ *p*_2_, the system evolves into stable patterned states (non-uniform biomass distributions). Each value of *p*_1_ and *p*_2_ was found as the average over 10 sets of simulations with different initial conditions and shortcuts positioning, the errorbars show the minimum and maximum values found within these 10 simulations.

To better illustrate this point, we study the dynamics of the patterned states on a precipitation gradient through continuation technique, for different *ϕ*_*b*_ values, similarly to what we did in figure 1 for the regular lattice case. Starting from the states reached at *p* = 1.7 through a disturbance of the uniform solution, we gradually change the precipitation while fixing the shortcut density and positions. At each step, we also add small amplitude uniformly distributed random noise to verify the stability of the solution. The results of these continuation series are shown in figure 5. We observe that the patterned states reached at *ϕ*_*b*_ ≳ 0.01 are more sensitive to precipitation changes and may be considered less resilient to environmental stress. This is in sharp contrast to the case of *ϕ*_*w*_ > 0 as we will see in section 3.2. At *ϕ*_*b*_ = 0 25, the uniform vegetation solution is stable through its whole existence range (*p* ≥ 1 52), and the system transitions through a tipping point to the desert state for *p* < 1.52.

**Figure 5:**
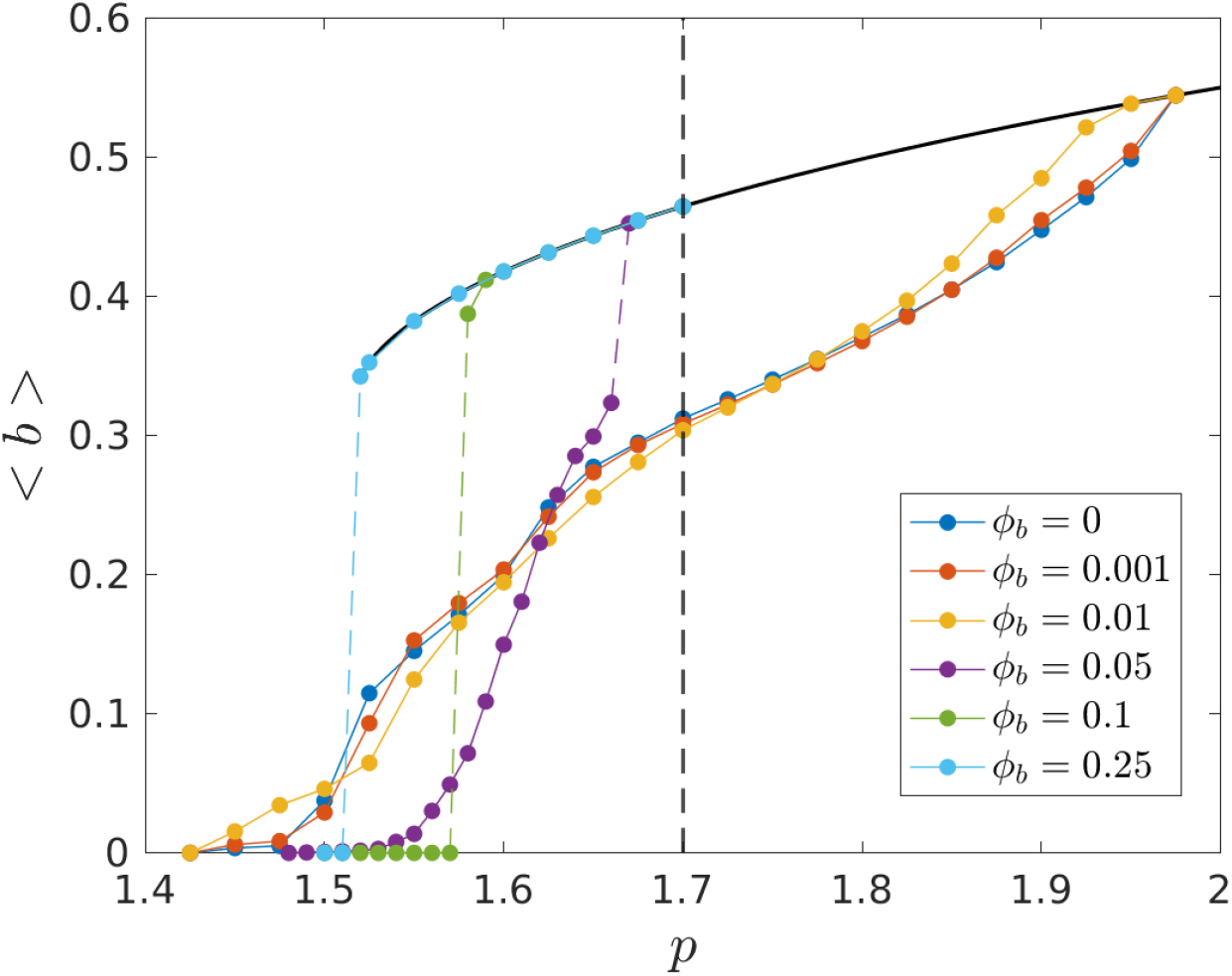
Results of five continuation series at different *ϕ*_*b*_ values, starting from the state reached at *p* = 1.7 from a disturbance of the uniform state. We modify the precipitation in small increments/decrements of 0.025. At each step (each precipitation change) we added uniformly distributed random noise in the interval [−0.05, 0.05] in order to verify the stability of the solution. The colored dashed line represent state transitions. The solid black line refers to the uniform vegetation solution, the dashed black line identifies the starting point of the continuation series.

We may deduce that, as the addition of randomly positioned shortcuts increases the connectivity of the biomass diffusion network, it facilitates the spread of biomass from vegetation-covered areas to the bare soil. Hence the system stabilizes around more spatially homogeneous patterns, up to the point where, at high shortcut densities, it loses the ability to form patterns entirely.

Further, as clearly visible in figure 3, at all precipitation values the patterns appear increasingly “disordered” for increasing *ϕ*_*b*_. The edges of the vegetated patches/bare gaps become frayed, and their shape and spacing is more heterogeneous. In other words, the addition of shortcuts reduces the regularity of the emerging patterns.

All patterns may be classified on a spectrum ranging from periodic to irregular. A periodic pattern is identical to itself when translated in space by some characteristic distance, while irregular patterns exhibit no identifiable periodicity or self-similarity at any spatial scale (Kästner et al., 2024b,a). Most natural patterns are regular but not periodic, meaning they become less self-similar when translated over increasingly larger distances (Kästner et al., 2024b). The regularity of a pattern is a measure of the distance over which the pattern loses its self-similarity.

In order to quantify pattern regularity in our simulations, we use the radial spectral density distributions (*S*_*r*_) (Kästner et al., 2024b,a). For each simulation, *S*_*r*_ is defined as the squared module of the 2-dimensional Fourier transform of the normalized biomass density distribution. For isotropic patterns such as dots, gaps, or labyrinths, we may convert the Fourier transform into polar coordinates 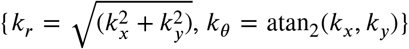 and integrate over the angular coordinate *k*_θ_ to obtain the radial spectral density as a function of . The radial spectral density distribution of a periodic pattern appears as an isolated peak, while regular non-periodic patterns show a lobed distribution which becomes wider for decreasing regularity (Kästner et al., 2024b).

Adopting the metric developed by Kästner et al. (2024b), we define the regularity index *R* of the patterns as the ratio between the peak of the radial spectral distribution (*S*_*rc*_ = max(*S*_*r*_)) and the characteristic wavelength (*λ*_*rc*_ |*S*_*r*_ (1/ *λ*_*rc*_) = *S*_*rc*_) . According to the meta-study conducted by the same authors (Kästner et al., 2024b), the median regularity of patterns generated through non-stochastic models is 2.36, while the median regularity of natural patterns is 0.67.

In figure 6, we show the radial spectral density of the patterns in figure 3, for *p* = 1.5 (dots pattern) and *p* = 1.7 (gaps pattern). We notice that, for increasing *ϕ*_*b*_, the central wavelength decreases in the dots pattern, as dots become more sparse, while it is roughly constant in the gaps pattern. All distributions become wider for increasing *ϕ*_*b*_. In the legend we report the regularity index at different *ϕ*_*b*_ values. The regularity of the patterns decreases for increasing *ϕ*_*b*_, growing closer to the median regularity of natural patterns (Kästner et al., 2024b).

**Figure 6:**
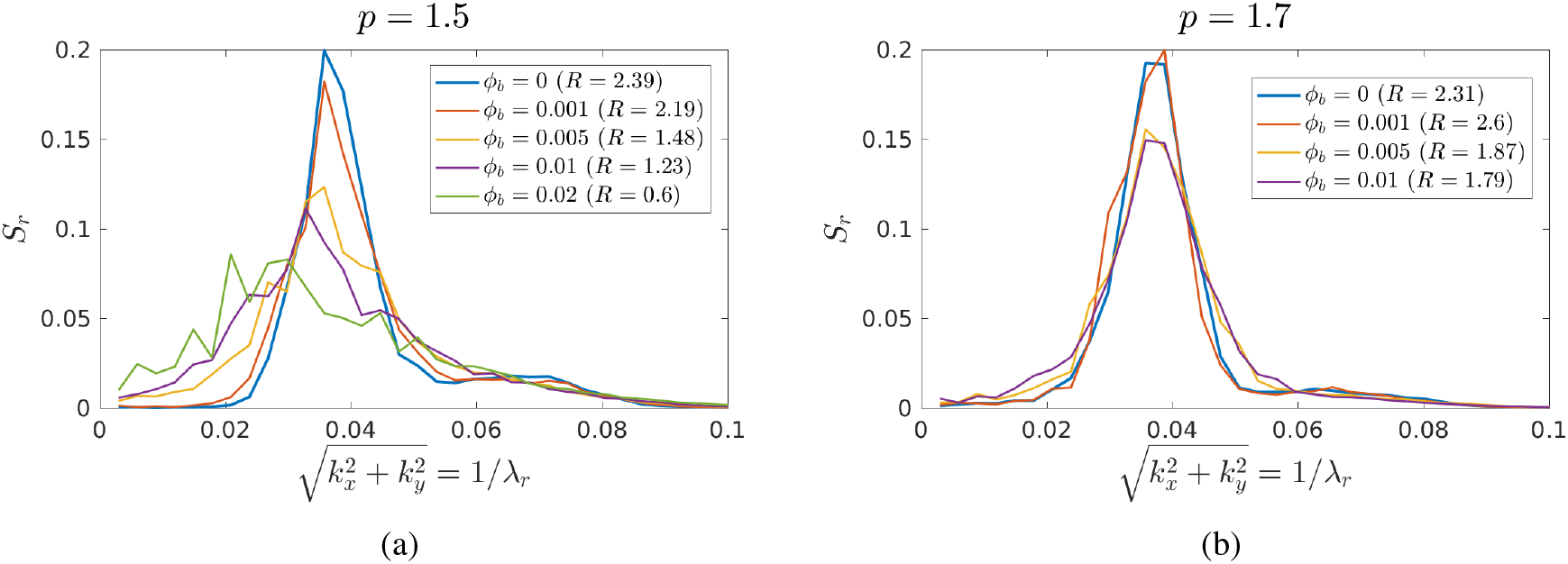
Radial spectral density distributions of the patterns shown in figure 3, at *p* = 1.5 (a) and *p* = 1.7 (b). The legend shows the values of *ϕ*_*b*_ and the corresponding regularity index *R*. Uniform states are not included in this figure.

However, at all *ϕ*_*b*_ values, the patterns preserve a characteristic wavelength and typical patch/gap size, differently, as we will see, from the results we obtained at *ϕ*_*w*_ > 0.

#### 3.1.1. Mean field variables

In addition to studying the spatial characteristics of the patterns, we analyze the behavior of a few spatially integrated variables with respect to *ϕ*_*b*_. This may help us gain a more intuitive understanding of the mechanisms behind the patterns in figure 3. Figure 7 shows the progression, for increasing *ϕ*_*b*_, of the average biomass density (7a), vegetation cover fraction (7b), average biomass density in vegetation-covered areas (7c), and average biomass density in vegetation-bare areas (7d). Each spatial point is classified as covered by vegetation if *b* > 0.22 (we choose this value to guarantee all non-uniform states include areas classified as bare).

**Figure 7:**
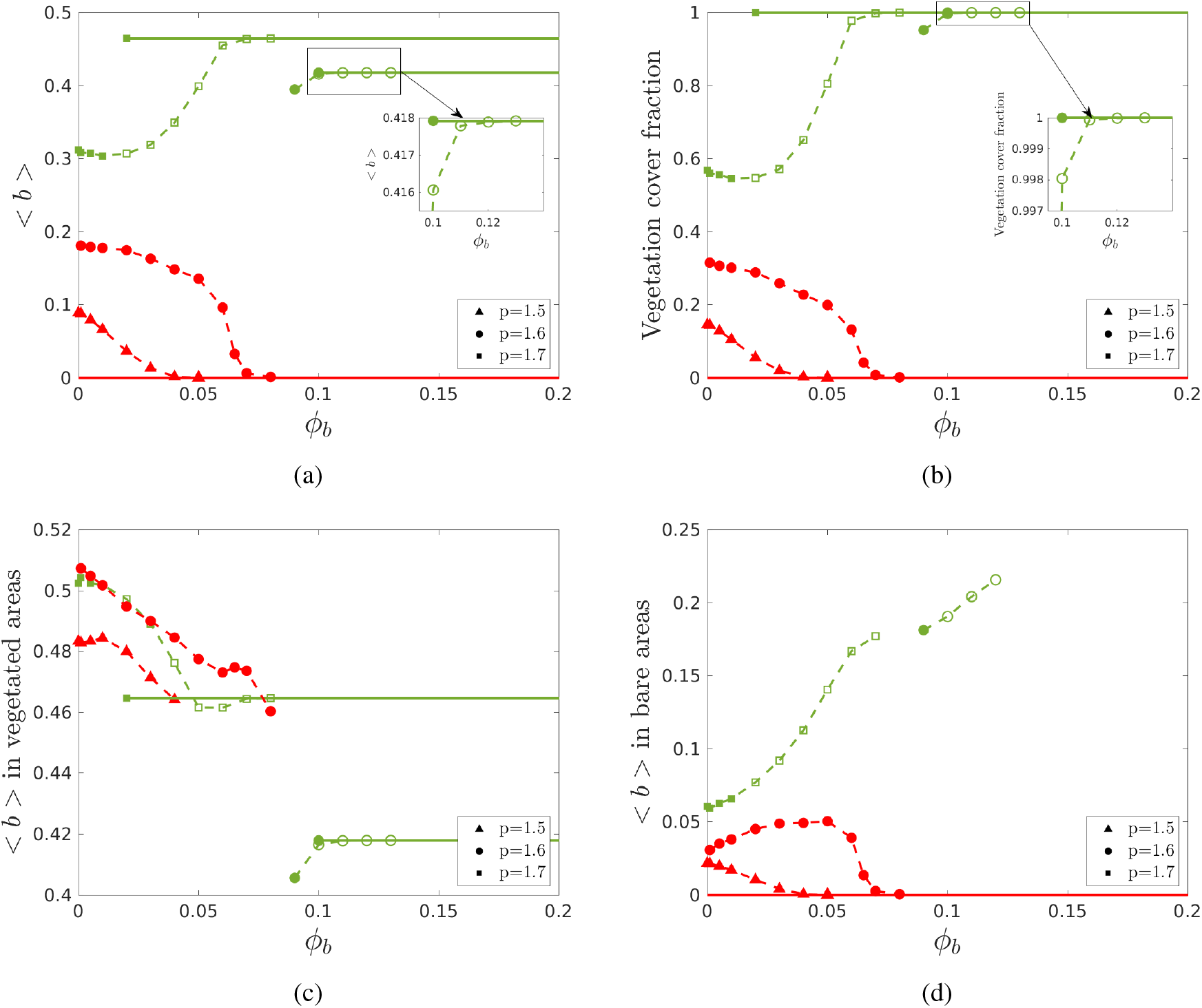
Spatially integrated variables in relation to *ϕ*_*b*_, for three different precipitation values. Green dashed lines refer to colonization patterns, red dashed lines refer to retreat patterns, solid lines refer to uniform states (green for uniform vegetation, red for desert). Each scatter point represents a simulation. Filled scatter points refer to solutions obtained starting from a uniform state perturbed with uniformly distributed noise in the range [-0.05, 0.05]. Non-filled scatter points refer to solutions obtained through continuation technique from the patterned states. The different plots show: average biomass density over the whole domain (a), fraction of the domain where *b* > 0.22 (b), average biomass density in vegetation-covered areas (*b* > 0.22) (c), and average biomass density in bare areas (*b* ≤ 0.22) (d).

As mentioned previously, the addition of shortcuts results in increased pressure towards spatial homogenization of the patterns. In different regions of the parameters space, the spatial homogenization can follow from two different processes. At sufficiently high precipitation/shortcut density values, the faster spread of biomass leads to the colonization of the vegetation-bare areas. This process produces low-regularity patterns which we termed *colonization patterns*, and, at sufficiently high shortcut density, it leads to the stability of the uniform vegetation solution. The dashed green lines in figure 7 show the progression of the colonization patterns towards the uniform vegetation solution. The insets in figure 7a and 7b serve to show the behavior of patterns very close to the uniform solution (such as the one shown in figure 3 at *p* = 1.6, *ϕ*_*b*_ = 0.09). The average biomass density and vegetation cover fraction of colonization patterns increase gradually with *ϕ*_*b*_, after an initial dip at low *ϕ*_*b*_ values (figure 7a-7b). The average biomass density within the vegetated areas decreases for increasing shortcut density until the patterned states merge with the uniform solution (figure 7c). Conversely, the biomass density of the bare areas increases steadily with *ϕ*_*b*_ as a result of the “colonization” process (figure 7d). At *p* = 1.7 colonization patterns are stable also at low *ϕ*_*b*_ values. At *p* = 1.6 colonization patterns are only stable for *ϕ*_*b*_ ≥ 0.09, hence the drastic shift in the patterns found from a disturbance of the uniform state (second row of figure 3).

At lower values of precipitation/shortcut density, the faster spread of biomass leads to the depletion of the vegetated patches, without the colonization of the vegetation-bare regions. Hence the system forms low-regularity patterns, here termed *retreat patterns*, whose biomass density and vegetation cover fraction decrease with *ϕ*_*b*_, until the desert solution becomes the only stable state. Red dashed lines in figure 7 show the progression of retreat patterns towards the desert state. As in colonization patterns, the biomass density of the vegetated areas decreases for increasing *ϕ*_*b*_. The biomass density of the bare regions, however, is also decreasing overall, though it may increase slightly at low values of *ϕ*_*b*_.

### 3.2. Water diffuses through a shortcut-augmented network (ϕ_b_ = 0, ϕ_w_ > 0)

We next study the impact of the structure of the water diffusion network by fixing *ϕ*_*b*_= 0 and increasing the parameter *ϕ*_*w*_. As we will see, diffusion of water over a network with intermediate shortcut density leads to the emergence of stable irregular patterns, meaning spatially heterogeneous states without a characteristic wavelength or a typical patch size. These are qualitatively different from the patterns analyzed in section 3.1, which showed lower regularity values than the patterns developing over a regular lattice, but could still be described through a characteristic wavelength and a typical patch size.

Figure 8 is equivalent to figure 3 and shows the final stage of 21 simulations, at 3 different precipitation values and 7 different *ϕ*_*w*_ values.

**Figure 8:**
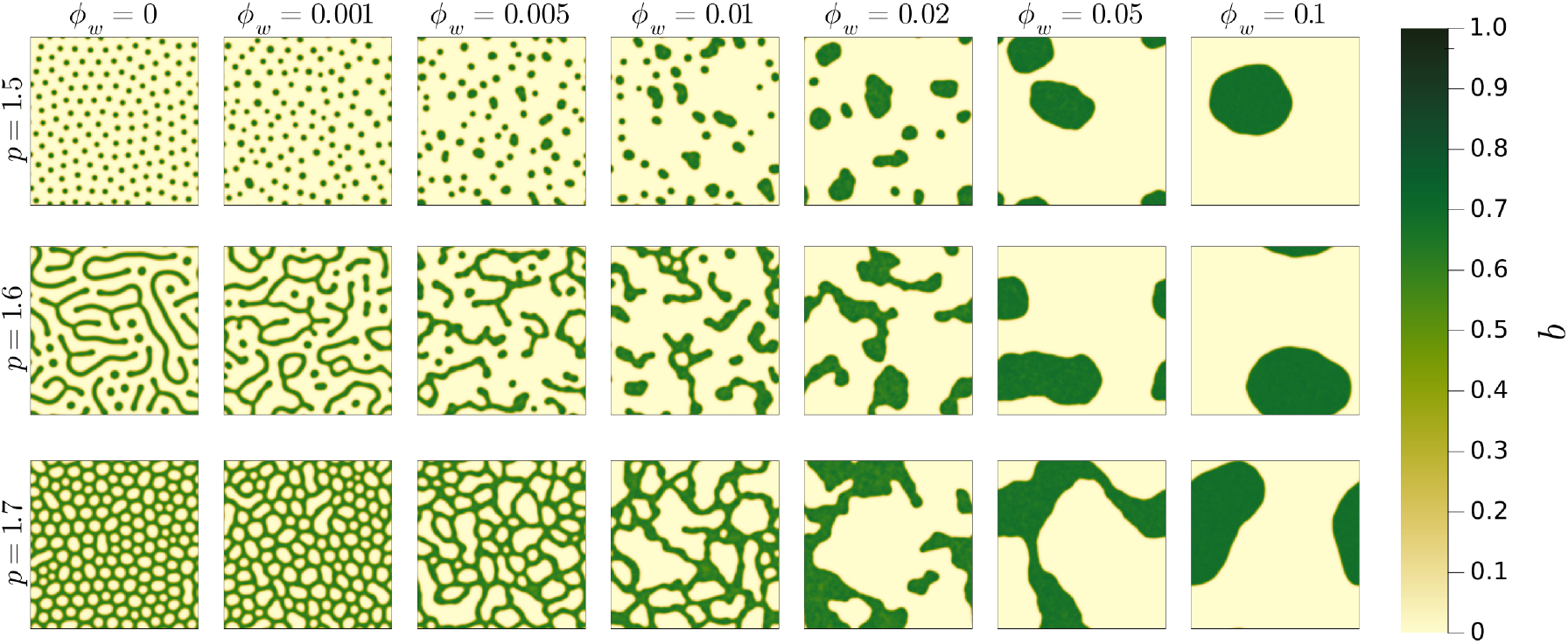
Biomass density distribution resulting from 21 different simulations at different *ϕ*_*w*_ and *p* values. The initial condition for all simulations is a uniform state perturbed with uniformly distributed random noise in the interval [−0.05, 0.05]. The simulation length is 5000 dimensionless times and the dimensionless domain size is 336×336 (400×400 points, spatial resolution 0.84).

As in the case discussed in section 3.1, the addition of shortcuts leads to a gradual decrease in the regularity of the emerging patterns. In this case, however, the loss of regularity is driven by a process of patch coarsening (Kletter et al., 2012), by which some vegetated patches/bare gaps increase in size while others shrink and disappear. At high shortcut density (*ϕ*_*w*_ ≳ 0.1), coarsening proceeds until all biomass in concentrated in a single, roughly circular patch over a bare landscape, with no qualitative difference between the three precipitation values. At lower values of *ϕ*_*w*_, the coarsening process slows down and eventually stops. In the intermediate shortcut density range (0.01 ≲ *ϕ*_*w*_ ≲ 0.02), the incomplete coarsening results in a final stage characterized by a broad distribution of patch and gap sizes, without a clear typical scale. In the Supplementary Material (S2), we represent the progression over time of the patch coarsening process.

Figure 9 shows the radial spectral density behavior for the patterns in figure 8, for *p* = 1 5 and *p* = 1 7. At all precipitation values, for increasing *ϕ*_*w*_, the spectral density behaviors become wider, and the patterns shift towards longer spatial wavelengths. At *ϕ*_*w*_ ≳ 0.01 the behaviors peak around the lowest non-zero frequency (characteristic wavelength is equal to or close to domain size). In the range 0.01 ≲ *ϕ*_*w*_ ≲ 0.02, this peak is a consequence of the irregularity of the patterns: we still observe numerous distinct patches/gaps within the domain (figure 3), but their distribution does not show any periodicity except that imposed by the periodic boundary conditions of the model (as opposed to the low-regularity patterns analyzed in figure 6, which showed a characteristic wavelength within the domain size). At higher *ϕ*_*w*_ values, all biomass is concentrated in a small number of similarly sized patches, or in a single patch for *ϕ*_*w*_ ≳ 0.1, so that the characteristic wavelength is naturally close to the domain size.

**Figure 9:**
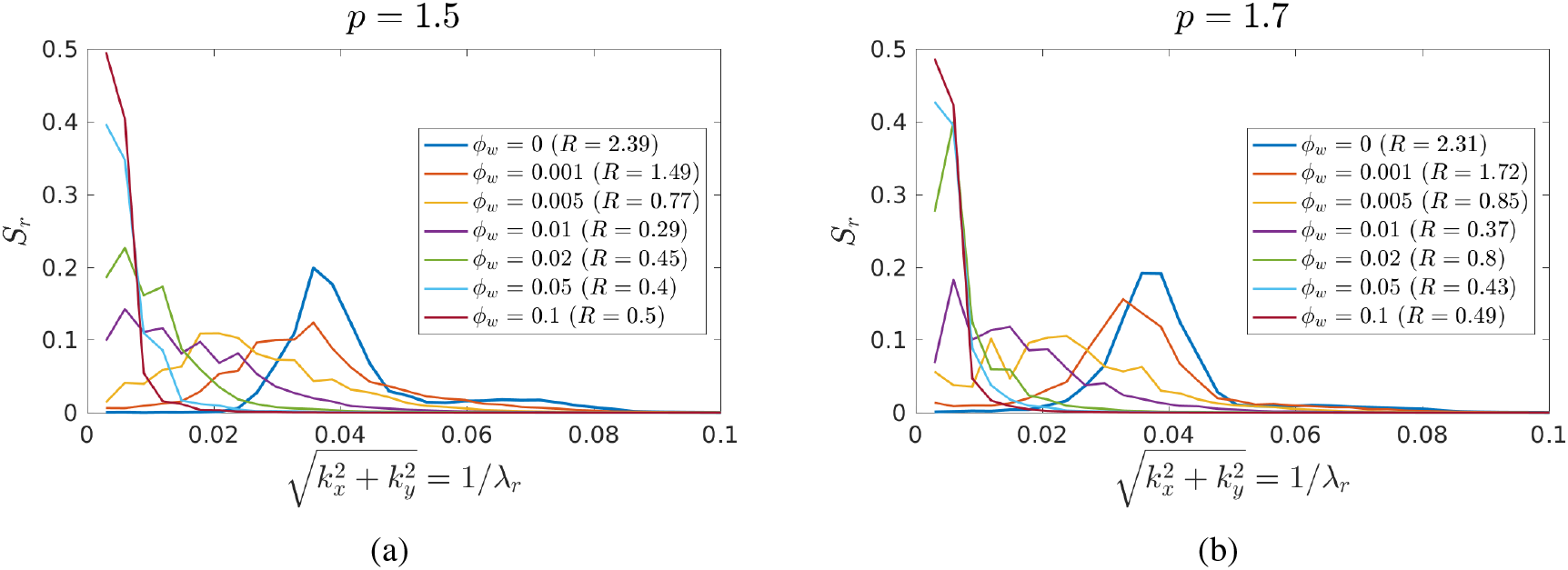
Radial spectral density behaviors of the patterns in figure 8, at *p* = 1.5 (a) and *p* = 1.7 (b). The legend shows the values of *ϕ*_*w*_ and the corresponding regularity index *R*.

Since the patterns observed in the intermediate *ϕ*_*w*_ range cannot be accurately described through their characteristic wavelength, or typical patch size, we aim at characterizing them through their patch size distributions. In figure 10 we show the exceedance distributions of patch sizes obtained at different *ϕ*_*w*_ values, for the case *p* = 1.5. A point was considered to be part of a patch if *b* > 0.05 (the choice of this threshold does not significantly affect the results). To improve the statistical robustness of the results, we computed the distributions over 25 ensemble runs for each *ϕ*_*w*_ value, using randomly positioned shortcuts at each run. We firstly notice that the range of patch sizes *s* increases for increasing *ϕ*_*w*_, as visible also in figure 8. We fit the exceedance distributions (by least mean square error method) with a truncated power law (*T P L*)

**Figure 10:**
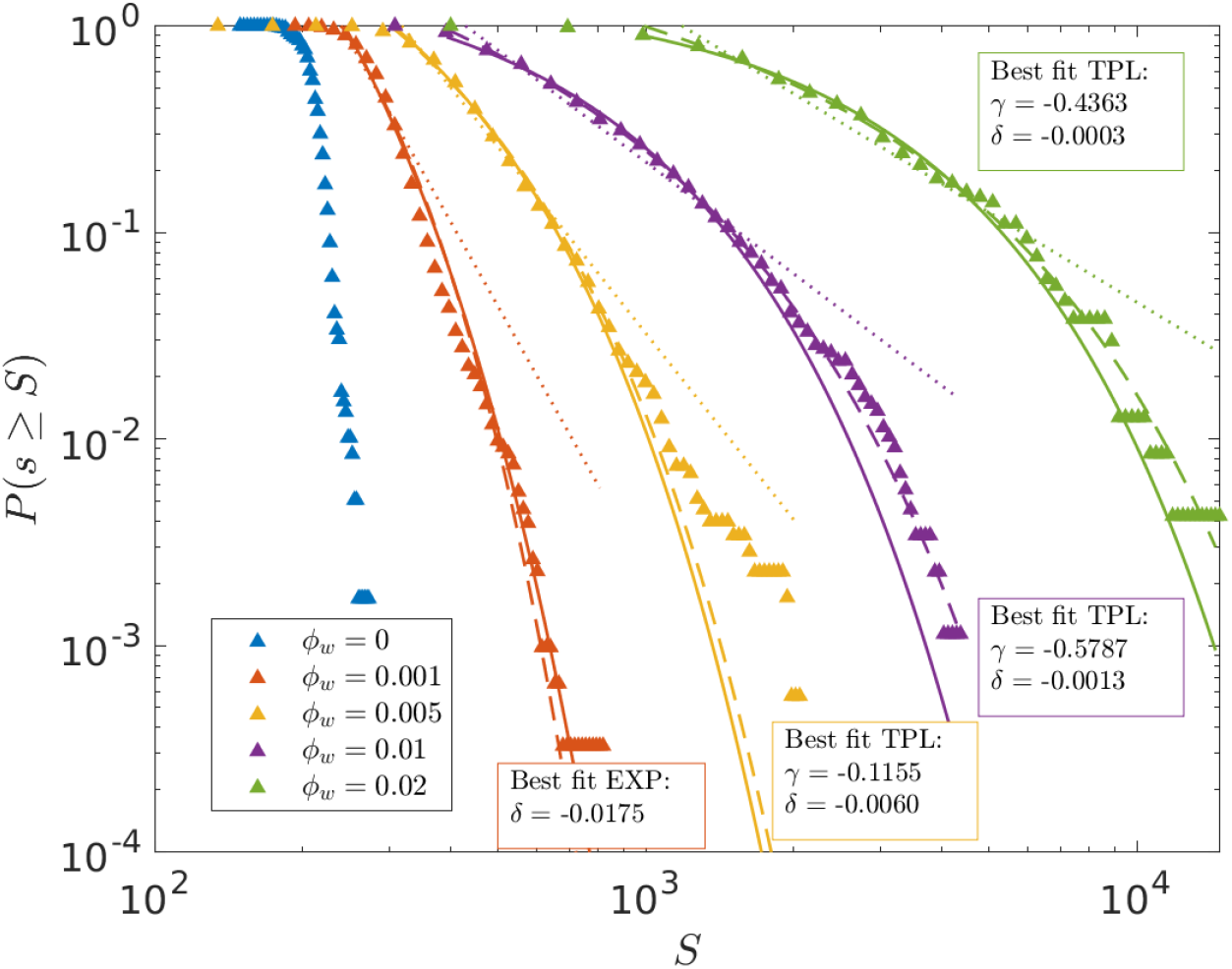
Exceedance distributions of vegetated patch sizes resulting from 25 ensemble simulations at *p* = 1.5, for different *ϕ*_*w*_ values. A point was considered to be part of a patch if *b* > 0.05. The distributions were fitted with the exponential law (solid line), the power law (dotted line) and the truncated power law (dashed line). We included in the fit only the largest 90% of patches.

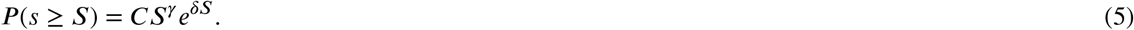

which reduces to an exponential function (EXP) when *γ* = 0, and a power law (PL) when *δ* = 0. If *γ* < 0 and *δ* < 0, then the inverse of the parameter *δ* determines the patch size above which the function decreases faster than a power law.

At *ϕ*_*w*_ = 10^−3^ our results were best fitted by the exponential function, at higher values of *ϕ*_*w*_ the truncated power law was the best fit. The parameter *δ* of the *T P L* decreases for increasing *ϕ*_*w*_, showing that the power law component of the distribution increases with shortcut density.

The irregular, wide patch size distribution patterns observed in the field often follow power laws or truncated power laws, and more rarely exponential functions (Kéfi et al., 2007; Lin et al., 2010; Maestre and Escudero, 2009; Scanlon et al., 2007). In section 4.3, we discuss these distributions in more details and explain their possible origin in our model.

Similarly to case of *ϕ*_*b*_ > 0 (section 3.1), the addition of shortcuts modifies not only the structure of the vegetation patterns, but also the precipitation range in which they emerge. Figure 11, equivalent to figure 4, shows the states emerging as a response to random disturbances of the uniform state, at different values of *p* and *ϕ*_*w*_. We notice that the lowest precipitation value in the stability range of the uniform vegetation solution (*p*_1_) increases for increasing *ϕ*_*w*_, while the lowest precipitation value at which patterns spontaneously emerge (*p*_2_) is constant in relation to *ϕ*_*w*_. In other words the addition of shortcuts to the water diffusion network expands the precipitation range for which the system self-organizes into patterns, contrary to the behavior observed in figure 4 over an augmented biomass diffusion network.

**Figure 11:**
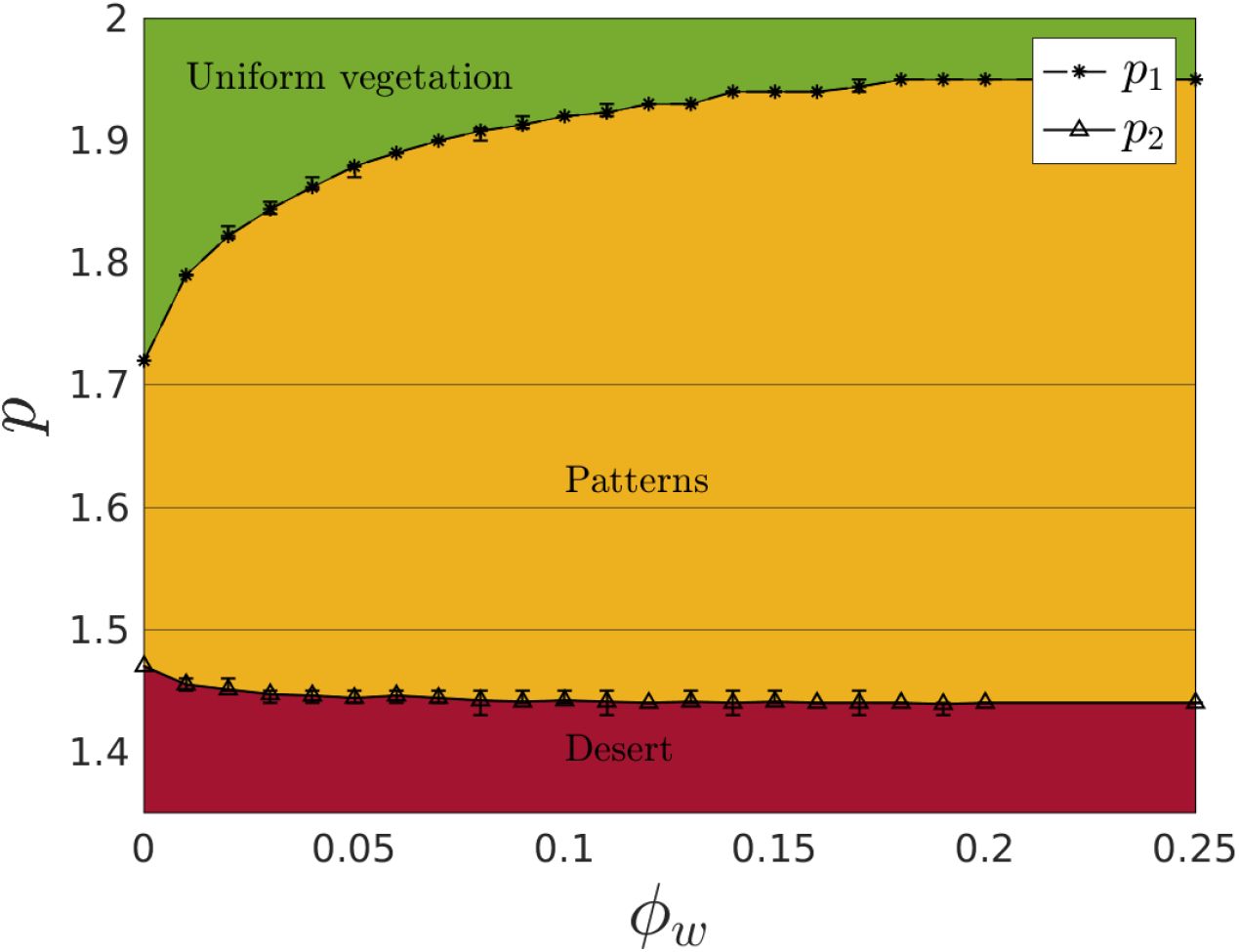
Diagram showing the final stage of simulations at different values of *p* and *ϕ*_*w*_, with initial condition uniform vegetation perturbed with random noise of maximum amplitude 0.05. The integration time and domain size are the same as in figure 8. For *p* ≥ *p*_1_ the uniform vegetation state is stable, and the perturbation is suppressed. For *p* < *p*_2_ the system evolves into a desert state. In the interval *p*_1_ > *p* ≥ *p*_2_, the system evolves into stable patterned states (non-uniform biomass distributions). Each value of *p*_1_ and *p*_2_ was found as the average over 10 sets of simulations with different initial conditions and shortcuts positioning, the errorbars show the minimum and maximum values found within these 10 simulations.

As pattern formation takes place over a wider precipitation range, we may expect the patterned states developing over an augmented water diffusion network, compared to the ones developing over a regular lattice, to exhibit a reduced sensitivity to precipitation changes and to persist on a wider range of precipitation values before merging with the uniform solutions. To strengthen these results, we repeat the experiment shown in figure 5, gradually changing the precipitation starting from the states found at *p* = 1.7, for different *ϕ*_*w*_ values. The results of these continuation series are shown in figure 12. As expected, for increasing *ϕ*_*w*_, the sensitivity of the patterned states to precipitation decreases, and patterned states persist at increasingly low/high precipitation values before merging with the desert/uniform vegetation state. Intuitively, this can be explained as a result of the more efficient exploitation of the water reserve by the vegetated patches due to the exploitation of fast subsurface transport mechanisms. Secondly, we observe the absence, at high *ϕ*_*w*_, of the steep slopes marking the transitions between different patterned states. As we add shortcuts, “dots”, “stripes” and “gaps” states gradually collapse into a single patterned state, establishing a linear relation between precipitation and average biomass density. This is already partly visible in figure 8, as the patterns generated at different precipitation values appear increasingly similar to each other for increasing *ϕ*_*w*_.

**Figure 12:**
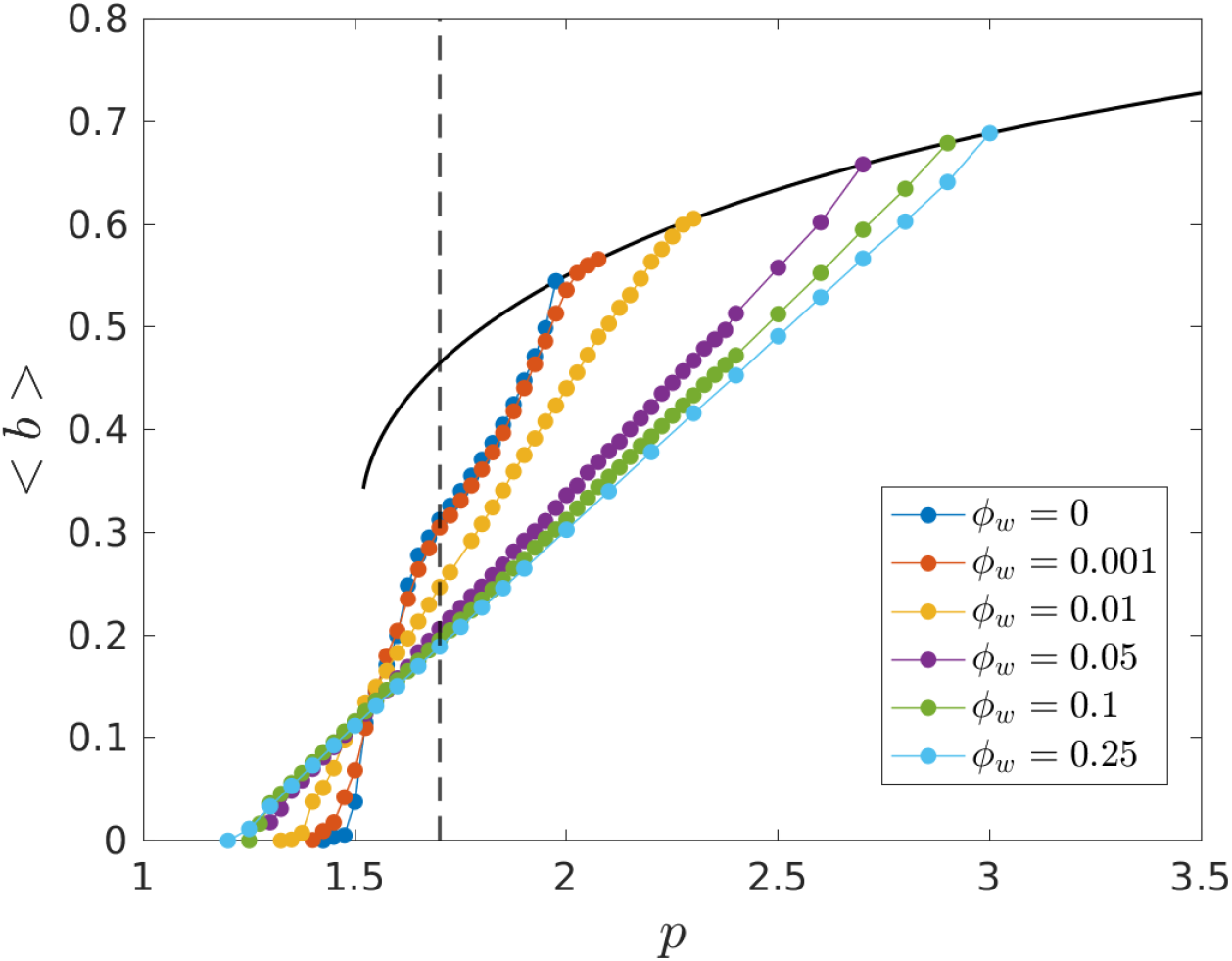
Results of five continuation series at different *ϕ*_*w*_ values, starting from the patterned state reached at *p* = 1.7 from a disturbance of the uniform state. The solid black line refers to the uniform vegetation solution, the black dashed line identifies the starting point of the continuation series.

#### 3.2.1. Mean field variables

As in section 3.1, we are interested in the behavior of some spatially integrated variables with respect to *ϕ*_*w*_, in order to gain a more intuitive understanding of the mechanisms at play. Figure 13 shows the progression, for increasing *ϕ*_*w*_, of the total average biomass density (13a), the vegetation cover fraction (13b), the average biomass density in vegetation-covered areas (13c), and the average biomass density in bare areas (13d). The three colors refer to three different precipitation values (*p* = 1.5, *p* = 1.6, *p* = 1.7).

**Figure 13:**
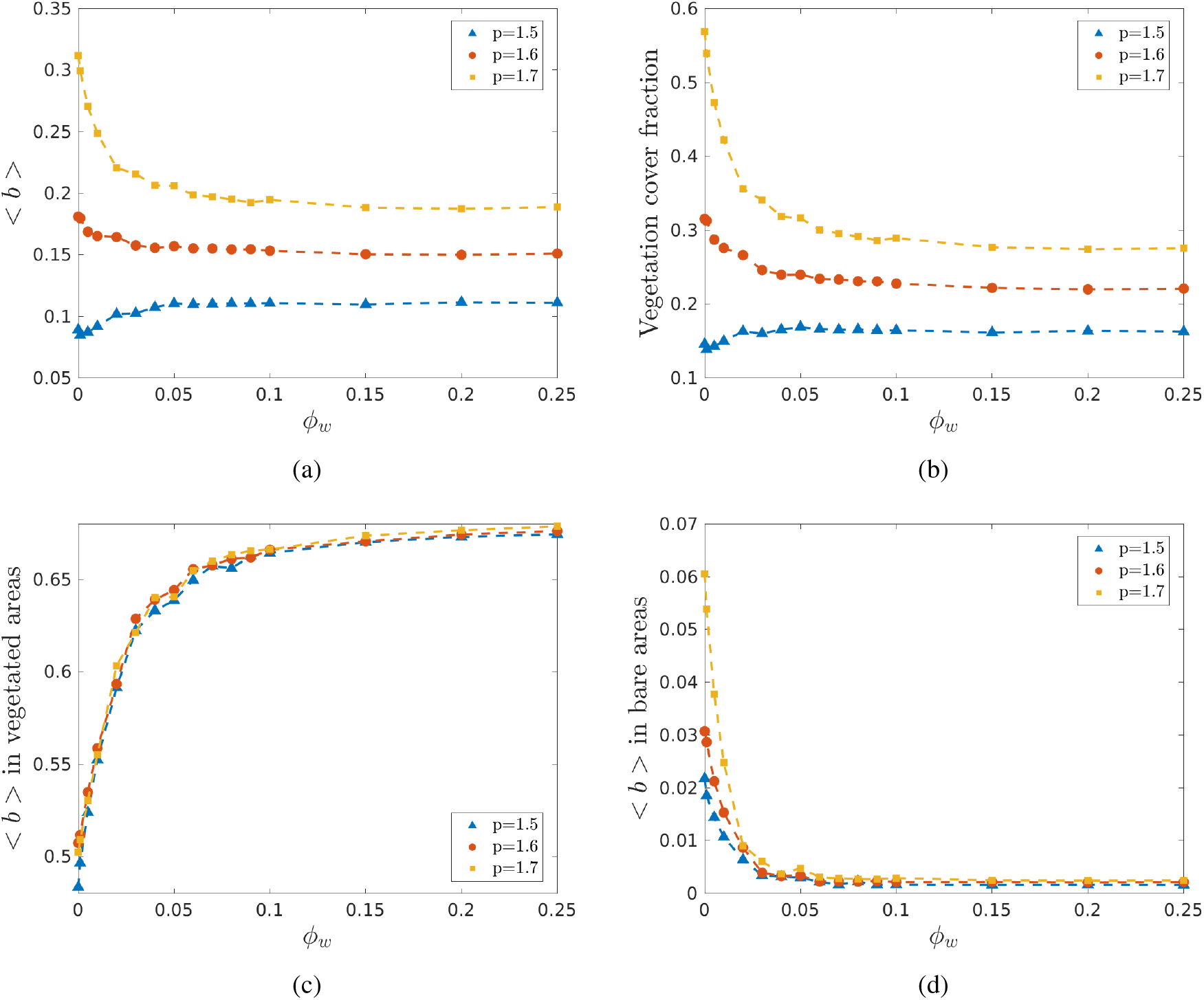
Behavior of the mean field variables in relation to *ϕ*_*w*_. Average biomass density over the whole domain (a), fraction of the domain where *b* > 0.22 (b), average biomass density in vegetation-covered areas (*b* > 0.22) (c), average biomass density in bare areas (*b* ≤ 0.22) (d).

The addition of shortcuts increases the connectivity of the water diffusion network and expands the area from which each patch is able extract water, leading to the growth in biomass density of the vegetated areas. As competition for the limited water resource takes place over a wider spatial range, some patches out-compete others, until the system reaches a new stable state characterized by a lower number of denser vegetation patches. As shown in figure 13c, the growth in biomass density within vegetated areas is the most visible effect of this process. Instead, biomass density in the bare areas decreases for increasing *ϕ*_*w*_ (figure 13d), as water is drained more thoroughly from the “water reserve” (the non-vegetated areas). Overall, while the addition of shortcuts to the biomass diffusion network leads to a spatial homogenization of the patterns (see section 3.1), the addition of shortcuts in the water diffusion network increases the “contrast” between vegetated and bare regions. Vegetation cover fraction (figure 13b) increases with *ϕ*_*w*_ at lower precipitation values (dots patterns) and decreases with *ϕ*_*w*_ at higher precipitation values (gaps and stripes patterns), as a result of the concentration of biomass into a lower number of continuous regions. The overall impact of the shortcuts on the average biomass density (figure 13a) depends on precipitation and it is the sum of two opposing effects: biomass growth in the vegetated areas and decrease in vegetation cover fraction. The patterns obtained at different precipitation values become closer to each other in terms of average biomass density, as the system becomes less sensitive to precipitation changes as already noted in figure 12. This behavior is in sharp contrast with what we observed in section 3.1 for an increase in *ϕ*_*b*_, where the shortcuts induced clearly different processes depending on precipitation.

## 4. Discussion

We have shown that the addition of long-distance transport links alters the pattern formation process in at least three key aspects: the precipitation range in which patterns emerge, the behavior of the spatially integrated variables, and the spatial structure of the patterns. The precipitation range in which we observe pattern formation shortens and eventually vanishes if biomass diffuses over an augmented network, while it expands if water diffuses over the same network. The addition of shortcuts to the biomass diffusion network leads to patterns characterized by a more spatially uniform distribution of biomass and a stronger sensitivity to precipitation in terms of their mean field variables. Conversely, shortcuts in the water diffusion network lead to patterns with a stronger contrast in biomass density between vegetated and bare areas, and lower sensitivity to precipitation. In terms of the impact on the spatial structure of the patterns, in both cases (*ϕ*_*b*_ > 0 and *ϕ*_*w*_ > 0) we may distinguish three blended ranges of shortcut densities. At lower densities, the regularity of the patterns does not appear significantly altered. At intermediate densities, we observe stable patterns characterized by low regularity (if biomass diffuses over an augmented network) or irregularity and broad patch size distributions (if water diffuses over an augmented network). At high densities, we detect either a uniform state or a patterned state characterized by a single patch over a bare landscape, when shortcuts are applied to the biomass or water diffusion network, respectively.

Some of these effects are easily explained as the result of increased overall biomass and water transport across the system. In order to single out the influence of spatial heterogeneity and multiple transport scales, we ran simulations of the local model (*ϕ*_*w*_ = *ϕ*_*b*_ = 0), varying the diffusion constants. The dimensionless diffusivity *d*_*w*_ in equations (1-2) represents the ratio of water and biomass diffusion constants, *D*_*W*_ /*D*_*B*_. Therefore, we compared the effects of an increase/decrease in *d*_*w*_, with the effects of diffusion over a shortcut-augmented water/biomass network. To this purpose we defined a parameter 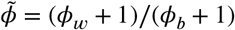, reflecting the ratio between the number of edges in the water and biomass diffusion networks. For the full discussion of this comparison we refer the reader to the Supplementary Material (S3), here we summarize the key findings.

We found that the qualitative behavior of the mean field variables in response to 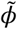 is coherent with changes in *d*_*w*_. Regarding the spatial structure of the patterns, we noticed that an increase in *d*_*w*_ leads to a homogeneous growth in patch size and characteristic wavelength, as well as an expansion of the precipitation range in which we observe patterns, as we found for 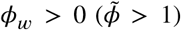. Conversely, a decrease in *d*_*w*_ leads to a decrease in patch size and the restriction of the pattern formation range, as we found for 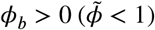. However, the low-regularity patterns found at intermediate *ϕ*_*b*_ values and the irregular, broad patch size patterns found at intermediate *ϕ*_*w*_ values do not emerge over a 2-dimensional lattice, at any values of diffusivity.

As mentioned in section 2.2, the addition of discrete long-distance transport links corresponds to the implementation of a Watts-Strogatz small world network model, whose properties are well-known within the field of network theory (Newman and Watts, 1999; Watts, 1999). In the following sections we apply network theory concepts in order to explain the pattern progression observed for increasing *ϕ*_*b*_ and *ϕ*_*w*_. In section 4.1 we consider the mean-field approximation of equations (3-4), valid in the high shortcut density limit. In section 4.2 we discuss the evolution of the topological properties of the network in comparison with the progression of pattern types. Finally, in section 4.3, we clarify the mechanism leading to the emergence of wide patch size distributions in the intermediate *ϕ*_*w*_ range.

### 4.1. Mean-field approximation

Dynamical systems operating on large random networks are commonly reduced to their mean-field approximation Barrat et al., 2008). This approximation scheme relies on the assumption that the state of each node is statistically independent from other network nodes, and rather depends on the average state of the system.

In our network model, we have a fixed grid component (the 2-dimensional lattice) and a growing random component (the shortcuts). In the limit of high number of shortcuts we may choose to represent the dynamics on the random part as their mean-field approximation. More specifically, in system (3-4), the adjacency matrices **A**^**b**^, **A**^**w**^ and the degree vectors **k**^**b**^, **k**^**w**^ are each the sum of a fixed lattice component 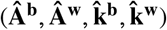 and a shortcut component, which depends on *ϕ*_*b*_ or 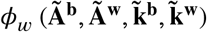. Preserving the representation of diffusion over the 2-dimensional lattice, for each node

*i*, we approximate the number of nodes connected to *i* through shortcuts (i.e. 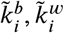) and the biomass and water density values of these nodes to their network averages. Hence we obtain the following mean-field model:

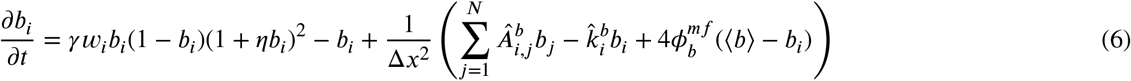

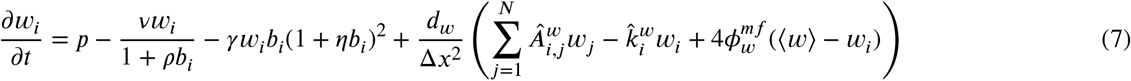

where ⟨.⟩ indicates spatial averaging 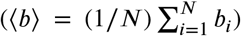, and we have redefined *ϕ*_*b*_ and *ϕ*_*w*_ as 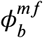 and 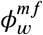, to distinguish between model (3-4) and model (6-7). Th^*i*=^e^1^full derivation of these equations from (3-4) can be found in the Supplementary Material (S4). We notice that, in this approximation, the representation of an heterogeneous environment disappears, and the influence of the added links is reduced a growing global coupling term, which relates the biomass and water densities of each point to their whole domain averages.

We first consider the case of 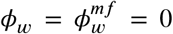. In figure 14, we show the biomass density maps resulting from simulations of the mean-field model (firstrow, 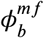 growing) and simulations of the full model (second row, *ϕ*_*b*_ growing). Considering the results of the mean-field model, we notice that at sufficiently low 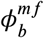 values (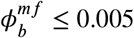 ≤ 0.005 at *p* = 1.5, 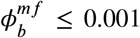 at *p* = 1.7), the patterned states persist, showing a decrease in the contrast between bare areas and vegetation covered areas, and the size of the vegetated dots at *p* = 1.5. This is similar to the behavior in the full model (second row), except the patterns preserve their regularity and characteristic wavelength. Beyond a certain 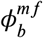 threshold, which depends on precipitation, the system transitions into either of the uniform solutions, reflecting the behavior of the full model at high *ϕ*_*b*_ .

**Figure 14:**
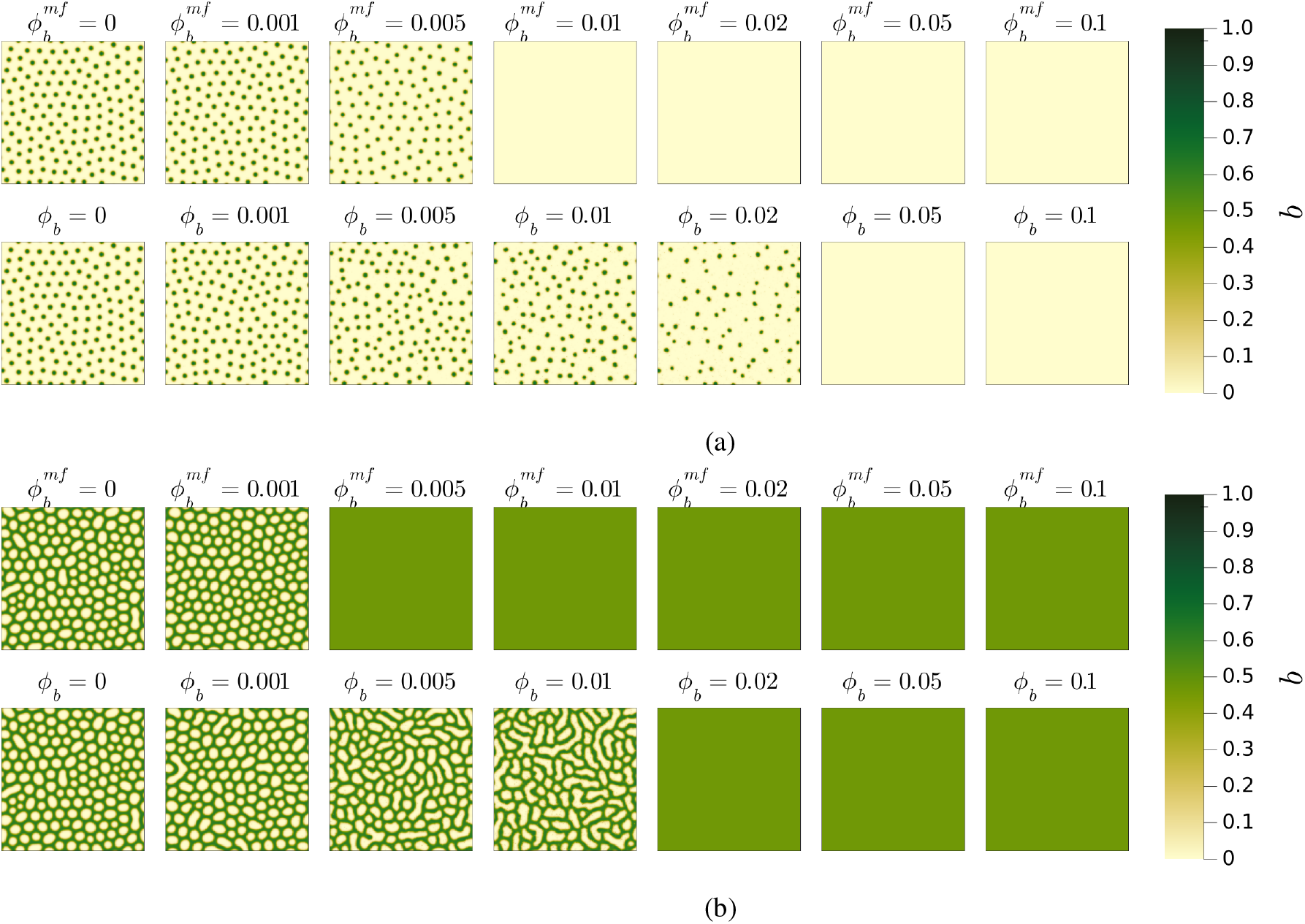
Comparison between the biomass density patterns obtained from the mean-field model (6-7) (first row) and from the full model (3-4) (second row), for 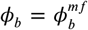 (increasing over the columns). In the two figures we consider two different precipitation values: *p* = 1.5 (a), *p* = 1.7 (b). The initial condition for all simulations is a uniform state perturbed with uniformly distributed random noise in the interval [−0.05, 0.05]. The simulation time was *t* = 5000. The domain size is 336×336 in dimensionless length.

Thus, the (6-7) approximation appears accurate at low *ϕ*_*b*_ values (*ϕ*_*b*_ ≲ 10^−3^) and high *ϕ*_*b*_ values (higher than the value beyond which we only find uniform states). However, in the intermediate range, the mean-field model is not able to reproduce the low-regularity patterns described in section 3.1 as retreat and colonization patterns. Rather, we find desert states instead of retreat patterns and uniform vegetation states instead of colonization patterns.

Figure 15 is equivalent to figure 14: here we consider the complementary case of the mean-field model for 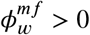 (first row) and the full model for *ϕ*_*w*_ > 0 (second row). Similarly to the case of 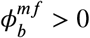, we notice a threshold behavior in the patterns produced through the mean-field model. At low intensities of the global coupling term, the system settles, over relatively short timescales (*t* ∼ 10^3^), on a regular pattern whose characteristic wavelength increases with 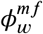, similarly to an increase in water diffusivity on a 2-dimensional lattice.

**Figure 15:**
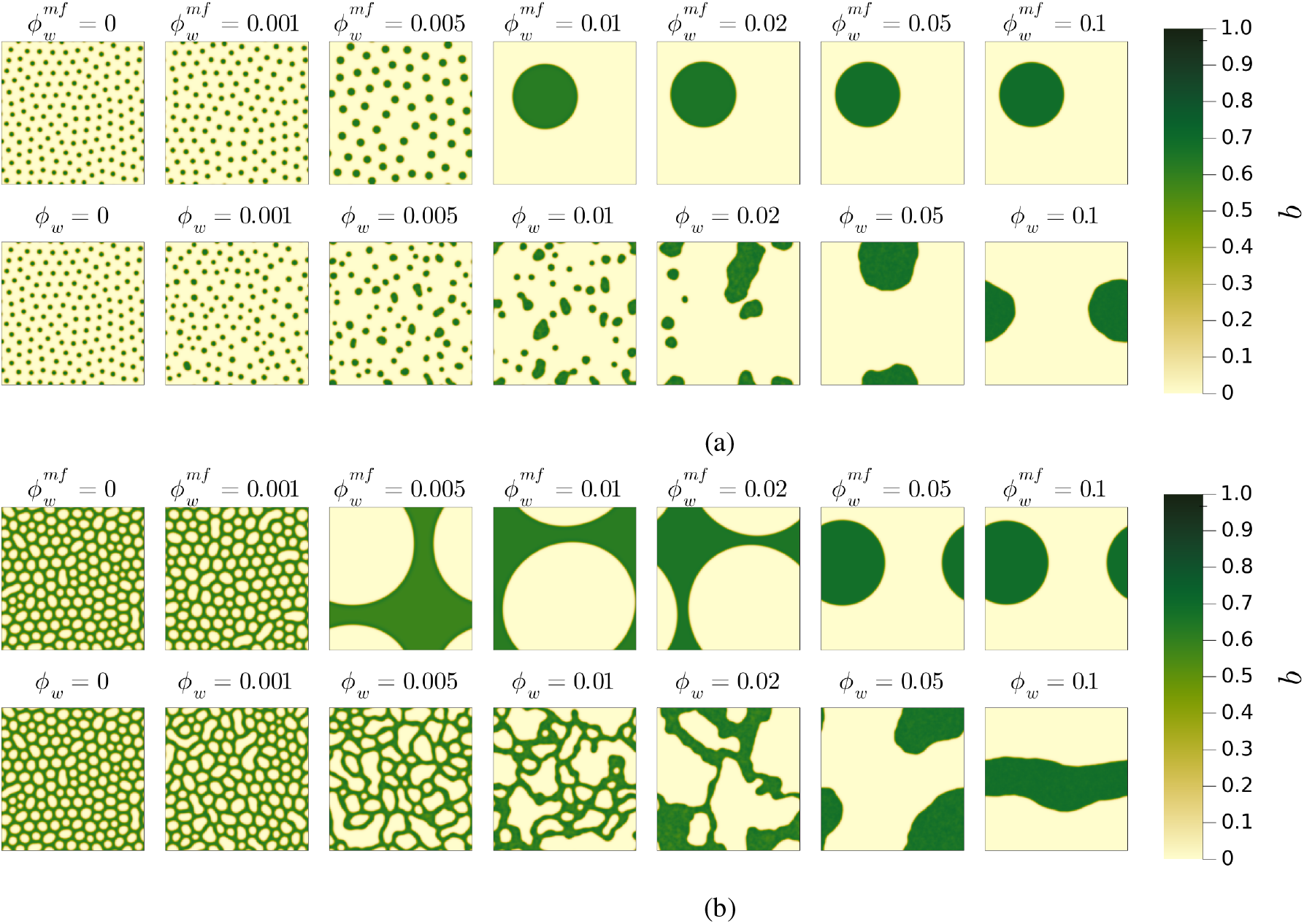
Comparison between the biomass density patterns obtained from the mean-field model (6-7) (first row) and from the full model (3-4) (second row), for 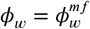 (increasing over the columns). In the two figures we consider two different precipitation values: *p* = 1.5 (a), *p* = 1.7 (b). The initial condition for all simulations is a uniform state perturbed with uniformly distributed random noise in the interval [−0.05, 0.05]. The simulation times were chosen in order to guarantee asymptotic stability. For 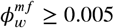, we chose longer simulation dimensionless times in the order of 10^4^ − 10^5^. The domain size is 336×336 in dimensionless length.

Above a certain threshold in 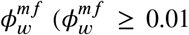 at *p* = 1.5, 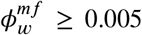 at *p* = 1.7), the patterns go through a very slow process of coarsening, in the course of which we observe transient states characterized by a wide patch size distribution. However, over very long timescales (*t* ∼ 10^4^ − 10^5^, simulation times in figure 15 were adjusted accordingly), we reach a state characterized by either a single circular patch over a bare landscape, or a single circular gap over a vegetation covered landscape. A single vegetation stripe over a bare landscape is also observed in some simulations. This behavior is similar to the one observed by Kletter et al. (2012) in a model subject to global competition for the water resource.

Intuitively, in model (6-7) growing 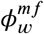 creates a pressure towards the homogenization of the water resource over the whole domain. Since the uniform solution is unstable, concentrating biomass in a continuous region allows the system to reduce water density in the bare regions, reducing the contrast with the vegetation covered regions. The coarsening process is initiated when the global pressure to homogenization wins over the local diffusion processes.

Thus, as in the case of *ϕ*_*b*_ > 0, at high shortcut density, model (6-7) is a good approximation of the full model, meaning the addition of shortcuts acts as a growing global coupling term, within this range. The irregular patterns found at intermediate densities, however, are not reproduced through the mean-field approximation. In these cases, the process of coarsening is initiated but it is interrupted as the patches become frozen into a broad patch size pattern.

We may also compare the patterns resulting from the two models in terms of their spatially averaged variables, as shown in the Supplementary Material (S5). We notice that, also in this case, the mean-field approximation is accurate at low 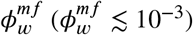 and at high 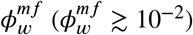, but deviation is larger in the intermediate range.

In conclusion, we can affirm that the model (6-7) undergoes a transition from a regime dominated by diffusion on the regular lattice to a regime dominated by the global coupling term, above certain 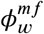 and 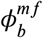 thresholds. This second regime represents a good approximation of the states reached at high shortcut densities in model (3-4). The behavior of model (6-7) may also be predicted analytically, as shown briefly in the Supplementary Material (S5).

### 4.2. Relation between network types and emerging patterns

The disordered patterns observed at intermediate values of *ϕ*_*b*_ and *ϕ*_*w*_ can not be reproduced through the mean-field approximation of the model, and are thus a consequence of the specific characteristics of the diffusion networks on which they develop. In this section, we discuss the patterns emerging at different values of *ϕ*_*b*_ and *ϕ*_*w*_ in comparison with the evolution of the topological properties of our network model.

In figure 16, we show the progression, for increasing *ϕ*, of two typical network metrics, on a system of fixed size (*N* = 400 × 400). These are the grid coefficient *c*_4_, and the average shortest path length *l*, both normalized for their value over a 2-dimensional lattice 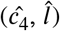. The grid coefficient *c*_4_ is an extension of the usual clustering coefficient *c*_3_ (which is always close to 0 in our model), and it refers to the relative abundance of quadrilaterals (groups of 4 nodes forming a cycle) over all sets of 4 connected nodes (Caldarelli et al., 2004; Imayama and Shiwa, 2008; Yin et al., 2018). In general, clustering coefficients indicate the tendency of a network to form “cliques” of densely connected nodes (Caldarelli et al., 2004). In our case, since the 2-dimensional lattice represents the real physical space on which patterns develop, the grid coefficient may be interpreted as a measure of the prevalence of local interactions. The average shortest path length is defined as the minimum number of edges to be crossed in order to connect two nodes of the network, averaged over the all pairs of nodes (Albert and Barabási, 2002; Watts and Strogatz, 1998). As an absolute value, it is often used to measure the efficiency of transport across a network (Watts and Strogatz, 1998). The scaling behavior of *l* with respect to the number of nodes *N* represents another important property of the network: *l* grows as 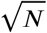 in a regular 2-dimensional lattice, while it grows as ln(*N*) or slower in random networks (Albert and Barabási, 2002; Fronczak et al., 2004). The insets in figure 16 show the relation between average path length and *N*, at four different *ϕ* values, together with a logarithmic fit.

**Figure 16:**
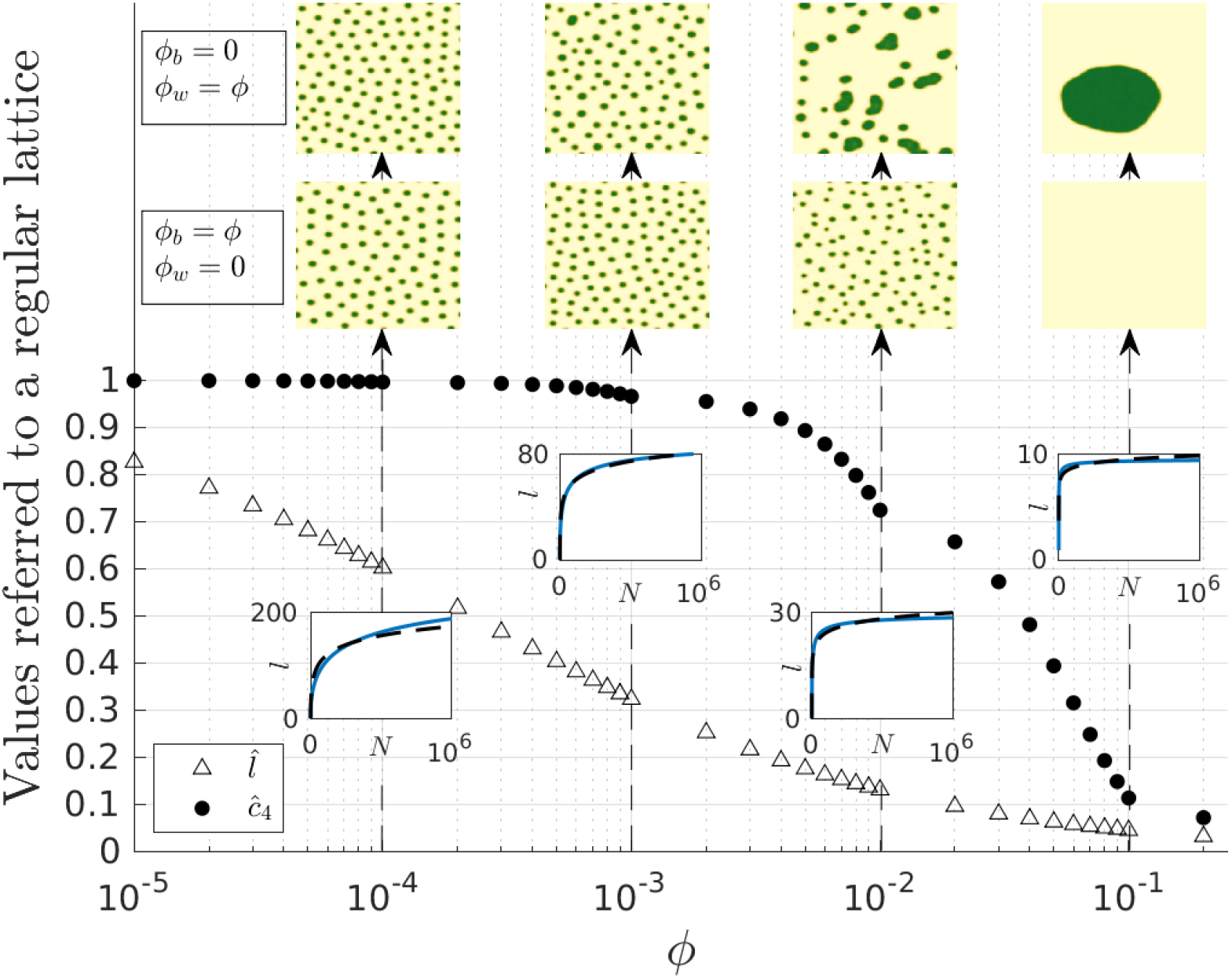
Behavior for increasing *ϕ* of the average shortest path length (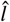, filled scatter points) and the grid coefficient (*ĉ*_4_, not filled scatter points) in our network model. Both variables are normalized by their value over a 2-dimensional lattice. The values of *ĉ*_4_ were calculated numerically, while 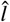 is based on the analytical approximation by Newman and Watts (1999). The insets show the behavior of the shortest path length *l* with system size (blue line) and a logarithmic fit (dashed black line), at 4 different *ϕ* values. The maps on top show the resulting patterns at 4 different *ϕ*_*b*_ and *ϕ*_*w*_ values.

From figure 16 (semilogarithmic in *ϕ*), we notice that, in the range *ϕ* = [0, 0.25], 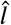 decreases close to linearly with log(*ϕ*), while *ĉ*_4_ starts to drop off at *ϕ* ∼ 10^−3^ (linear decrease with *ϕ*). From the insets, we verify that the scaling of *l* with system size transitions gradually from quadratic to logarithmic, with the crossover at *ϕ* ∼ 10^−3^, as expected from previous studies (Barrat and Weigt, 2000).

Thus, we may distinguish three types of networks generated by our model based on their topological properties. Each type, when applied to the biomass or water diffusion networks of our vegetation model, leads to the formation of different patterns in the real space, as shown in the maps on top of figure 16. At low shortcut density (approximately *ϕ* < 10^−3^), the grid coefficient is almost unchanged from that of a 2-dimensional lattice. The average shortest path length is significantly reduced, but it grows faster than logarithmically with system size (first inset at *ϕ* = 10^−4^). The patterns developing in this range are similar to the ones observed over a 2-dimensional lattice in terms of their spatial structure. The improved efficiency of transport through the reduced shortest path length affects only the mean field variables of the patterns, whose behavior is consistent with a change in the diffusion constants over a 2-dimensional lattice (see Supplementary Material (S3)). In the intermediate *ϕ* range (approximately 10^−3^ < *ϕ* ≤ 5 ⋅ 10^−2^), we have still a high grid coefficient, while the average shortest path length is much lower than that of a regular lattice, and it scales logarithmically or slower than logarithmically with system size (insets at *ϕ* = 10^−3^ and *ϕ* = 10^−2^). This type of network (high clustering, low average path length) can be described as small world (Barrat and Weigt, 2000; Watts and Strogatz, 1998). A wide variety of real-world networks show this kind of topology (Bassett and Bullmore, 2017; Olesen et al., 2006; Telesford et al., 2011). In this range we observe low-regularity vegetation patterns with a typical patch size (if biomass diffuses through the small world network), or irregular broad patch size patterns (if water diffuses through the small world network). Finally, at high shortcut density (approximately *ϕ* > 5 ⋅ 10^−2^) the topology resembles a random network, with very low *l* and *c*_4_ compared to a regular lattice, and slower than logarithmic scaling of *l* with system size (inset at *ϕ* = 10^−1^). In this range, the mean-field approximation of the model becomes valid, and we observe either uniform states (if biomass diffuses through the random network), or single patch/gap states (if water diffuses through the random network).

Thus, the disordered patterns found in the small world range may be interpreted as intermediate “frozen” phases between the patterns regulated by local diffusion and the ones governed by global coupling. At *p* = 1.5, when biomass operates over a small world network, some vegetated patches shrink and others disappear as a result of diffusion of biomass onto the bare soil, but the process becomes frozen in time before the desert state is reached. Similarly, when water operates over a small world network, the broad patch size patterns are the result of a frozen coarsening process.

### 4.3. The origin of wide patch size distributions at intermediate shortcut densities

To illustrate the mechanism driving the formation of disordered patterns at intermediate shortcut densities, we focus on the wide patch size distribution that emerges at intermediate *ϕ*_*w*_.

To this purpose, we introduce a different topological property of a network: the size, *V* (*r*), of a neighborhood of radius *r*. That is defined as the number of nodes reachable through a maximum of *r* edges from a given node, averaged over all nodes of the network. Intuitively, for a network generated through our model at a certain value of *ϕ, V* (*r, ϕ*) is some growing function of *r* and *ϕ*. More quantitatively, Newman and Watts (1999) found that, in a network model like the one adopted here, *V* (*r, ϕ*) is related to *ϕ* through the characteristic length of the network 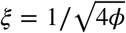, which refers to the typical distance between the ends of two separate shortcuts. Newman and Watts (1999) showed that *V* (*r, ξ*) grows proportionally to *r*^2^ if *r* ≪ *ξ*, while it grows as *e*^*r*/*ξ*^ if *r* ≫ *ξ*.

Focusing on the case of water diffusing over a shortcut-augmented network (*ϕ* = *ϕ*_*w*_), and considering a vegetated patch of size in the physical space, we can define the characteristic length of the patch as 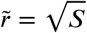. The size of this same patch when mapped onto the water diffusion network – meaning the number of points from which the patch can draw water – is proportional to 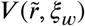, with 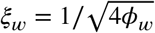.

Figure 17 shows the behavior for increasing *ϕ*_*w*_ of the characteristic length *ξ*_*w*_ of the network, as well as the smallest, average and largest characteristic lengths of the vegetated patches in the emerging patterns 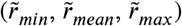. As we cross in the small world range (*ϕ* > 10^−3^), the characteristic lengths of the patches become larger than the characteristic length of the network *ξ*_*w*_, and an exponential relation is established between the size of the patch and the number of points from which it may draw water. Thus larger patches are exponentially favored to expand and out-compete smaller ones, leading to the coarsening process previously described. However, as long as the network is highly clustered (*ϕ* ≤ 5 ⋅ 10^−2^), and the dynamics can not be reduced to their mean-field approximation, the coarsening process is slow enough that global resources are exhausted before a single patch is able to out compete all others. Hence the system becomes stuck in a broad patch size distribution pattern, as shown in figure 10.

**Figure 17:**
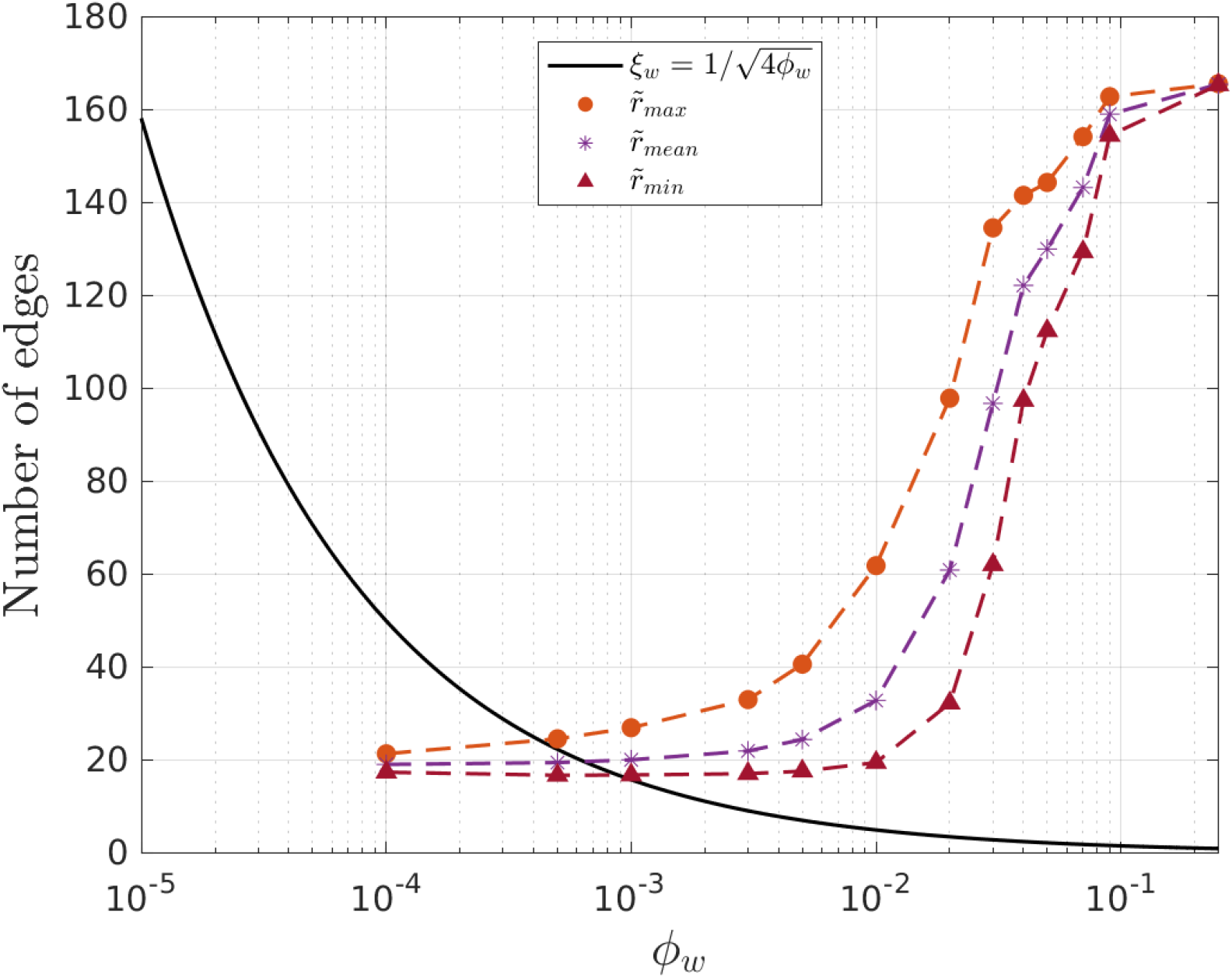
The solid black line shows the behavior for increasing *ϕ*_*w*_ of the characteristic length of the water diffusion network *ξ*_*w*_. The scatter points show the characteristic length 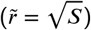 of the largest, average and smallest patch of the resulting patterns at *p* = 1.5. These values are averaged over 25 simulations with different shortcut positions.

In physical terms, the availability of domain scale water transport mechanisms produces a scale-dependent feedback that favors the growth of larger patches and leads to coarsening, as in previous studies focusing on global competition (von Hardenberg et al., 2010; Kletter et al., 2012). However, in our model, the sparseness and spatial heterogeneity of access to these alternative transport mechanisms stabilizes the transient broad patch size distribution patterns.

Obviously, a similar mechanism is active in the case of biomass diffusion over a shortcut-augmented network. However, since the faster diffusion of biomass tends to disperse rather than expand the largest patches, patch sizes never cross over into the exponential growth region, producing patterns with decreasing regularity but no wide patch size distribution.

## 5. Conclusion

Based on previous studies exploring the effects of spatial heterogeneity and domain scale transport on dryland vegetation patterns (von Hardenberg et al., 2010; Pinto-Ramos et al., 2025; Yizhaq et al., 2014), we have proposed a new approach to represent the spatially heterogeneous co-existence of different transport scales in a reaction-diffusion vegetation model. In our model the local availability of domain scale transport mechanisms is represented through the addition of randomly distributed shortcuts over a 2-dimensional lattice. This method has the advantage of a negligible increase in computational cost compared to the local model, and the possibility to exploit the direct analogue with network theory.

In an intermediate shortcut density range producing a small world network topology, we observed two distinct kinds of stable disordered patterns. Low-regularity patterns emerged when sparse shortcuts were applied to the biomass diffusion network, while irregular patterns with a broad patch size distribution appeared when they were applied to the water diffusion network. This range corresponds to systems in which domain scale transport is possible, but only sparsely available, based on local conditions. Crucially, these two kinds of disordered patterns arise from different mechanisms, and show different behaviors for varying precipitation. Compared to regular patterns, low-regularity patterns showed a decreased resilience to environmental changes, while irregular patterns showed an increased resilience.

In section 2.2 we have discussed examples of transport mechanisms that fit our model description. Nonetheless, our model is quite idealized and would likely require some adjustments to be applied to any specific environment: for instance, in the form of a time-varying diffusion network, or a distance-decaying kernel applied to the positioning of shortcuts. The idealization, however, allows us to expand the applicability of our results to a wide range of other environments subject to similar pattern-forming mechanisms (Rietkerk and van De Koppel, 2008).

## CRediT authorship contribution statement

**Sara Filippini:** Conceptualization, Methodology, Software, Formal Analysis, Investigation, Data Curation, Writing-Original Draft, Visualization. **Luca Ridolfi:** Conceptualization, Methodology, Resources, Writing-Reviewing and Editing, Supervision, Project Administration, Funding acquisition. **Jost von Hardenberg:** Conceptualization, Methodology, Software, Resources, Writing-Reviewing and Editing, Supervision, Project Administration, Funding acquisition.

## Data availability

The computational model used in this study is archived and publicly available at 10.5281/zenodo.18132089

## References

Albert, R., Barabási, A.L. 2002. Statistical mechanics of complex networks. Reviews of Modern Physics 74, 47–97. doi:10.1103/RevModPhys.74.47.

Band, L.E., McDonnell, J.J., Duncan, J.M., Barros, A., Bejan, A., Burt, T., Dietrich, W.E., Emanuel, R.E., Hwang, T., Katul, G., Kim, Y., McGlynn, B., Miles, B., Porporato, A., Scaife, C., Troch, P.A. 2014. Ecohydrological flow networks in the subsurface. Ecohydrology 7, 1073–1078. doi:10.1002/eco.1525.

Barrat, A., Barthélemy, M., Vespignani, A. 2008. Dynamical Processes on Complex Networks. Cambridge University Press, Cambridge.doi:10.1017/CBO9780511791383.

Barrat, A., Weigt, M. 2000. On the properties of small-world network models. The European Physical Journal B - Condensed Matter and Complex Systems 13, 547–560. doi:10.1007/s100510050067.

Bassett, D.S., Bullmore, E.T. 2017. Small-World Brain Networks Revisited. The Neuroscientist 23, 499–516. doi:10.1177/1073858416667720.

Behrend, D., Athmann, M., Han, E., Küpper, P.M., Perkons, U., Bauke, S.L., Köpke, U., Kautz, T., Gaiser, T., Seidel, S.J. 2025. Do biopores created by perennial fodder crops improve the growth of subsequent annual crops? A synthesis of multiple field experiments. Field Crops Research 322, 109687. doi:10.1016/j.fcr.2024.109687.

Borgogno, F., D’Odorico, P., Laio, F., Ridolfi, L. 2009. Mathematical models of vegetation pattern formation in ecohydrology. Reviews of Geophysics 47, RG1005. doi:10.1029/2007RG000256.

Bullock, J.M., Clarke, R.T. 2000. Long distance seed dispersal by wind: measuring and modelling the tail of the curve. Oecologia 124, 506–521.doi:10.1007/PL00008876.

Buttle, J.M., McDonald, D.J. 2002. Coupled vertical and lateral preferential flow on a forested slope. Water Resources Research 38.doi:10.1029/2001WR000773.

Caldarelli, G., Pastor-Satorras, R., Vespignani, A. 2004. Structure of cycles and local ordering in complex networks. The European Physical Journal B - Condensed Matter 38, 183–186. doi:10.1140/epjb/e2004-00020-6.

Cross, M.C., Hohenberg, P.C. 1993. Pattern formation outside of equilibrium. Reviews of Modern Physics 65, 851–1112. doi:10.1103/RevModPhys.65.851.

Dijkstra, H.A. 2011. Vegetation Pattern Formation in a Semi-Arid Climate. International Journal of Bifurcation and Chaos 21, 3497–3509. doi:10.1142/S0218127411030696.

D’Odorico, P., Laio, F., Ridolfi, L. 2006. Patterns as indicators of productivity enhancement by facilitation and competition in dryland vegetation.Journal of Geophysical Research: Biogeosciences 111, G03010. doi:10.1029/2006JG000176.

Eigentler, L., Sherratt, J.A. 2018. Analysis of a model for banded vegetation patterns in semi-arid environments with nonlocal dispersal. Journal of Mathematical Biology 77, 739–763. doi:10.1007/s00285-018-1233-y.

Freschet, G.T., Pagès, L., Iversen, C.M., Comas, L.H., Rewald, B., Roumet, C., Klimešová, J., Zadworny, M., Poorter, H., Postma, J.A., Adams, T.S.,Bagniewska-Zadworna, A., Bengough, A.G., Blancaflor, E.B., Brunner, I., Cornelissen, J.H.C., Garnier, E., Gessler, A., Hobbie, S.E., Meier, I.C., Mommer, L., Picon-Cochard, C., Rose, L., Ryser, P., Scherer-Lorenzen, M., Soudzilovskaia, N.A., Stokes, A., Sun, T., Valverde-Barrantes, O.J., Weemstra, M., Weigelt, A., Wurzburger, N., York, L.M., Batterman, S.A., Gomes De Moraes, M., Janeček, S., Lambers, H., Salmon, V., Tharayil, N., McCormack, M.L. 2021. A starting guide to root ecology: strengthening ecological concepts and standardising root classification, sampling, processing and trait measurements. New Phytologist 232, 973–1122. doi:10.1111/nph.17572.

Fronczak, A., Fronczak, P., Hołyst, J.A. 2004. Average path length in random networks. Physical Review E 70, 056110. doi:10.1103/PhysRevE.70.056110.

Fukui, S., Araki, K.S. 2014. Spatial Niche Facilitates Clonal Reproduction in Seed Plants under Temporal Disturbance. PLoS ONE 9, e116111. doi:10.1371/journal.pone.0116111.

Gilad, E., von Hardenberg, J., Provenzale, A., Shachak, M., Meron, E. 2004. Ecosystem Engineers: From Pattern Formation to Habitat Creation.Physical Review Letters 93, 098105. doi:10.1103/PhysRevLett.93.098105.

Graham, C.B., Lin, H. 2012. Subsurface Flow Networks at the Hillslope Scale, in: Hydropedology. Elsevier, pp. 559–593. doi:10.1016/B978-0-12-386941-8.00018-6.

Halatek, J., Brauns, F., Frey, E. 2018. Self-organization principles of intracellular pattern formation. Philosophical Transactions of the Royal Society B: Biological Sciences 373, 20170107. doi:10.1098/rstb.2017.0107.

von Hardenberg, J., Kletter, A.Y., Yizhaq, H., Nathan, J., Meron, E. 2010. Periodic versus scale-free patterns in dryland vegetation. Proceedings of the Royal Society B: Biological Sciences 277, 1771–1776. doi:10.1098/rspb.2009.2208.

von Hardenberg, J., Meron, E., Shachak, M., Zarmi, Y. 2001. Diversity of Vegetation Patterns and Desertification. Physical Review Letters 87, 198101. doi:10.1103/PhysRevLett.87.198101.

Hardie, M.A., Doyle, R.B., Cotching, W.E., Lisson, S. 2012. Subsurface Lateral Flow in Texture-Contrast (Duplex) Soils and Catchments with Shallow Bedrock. Applied and Environmental Soil Science 2012, 861358. doi:10.1155/2012/861358.

Hartnett, D.C., Setshogo, M.P., Dalgleish, H.J. 2006. Bud banks of perennial savanna grasses in Botswana. African Journal of Ecology 44, 256–263. doi:10.1111/j.1365-2028.2006.00646.x.

Imayama, R., Shiwa, Y. 2008. Lamellar pattern formation in small-world media. Physica A: Statistical Mechanics and its Applications 387,1033–1048. doi:10.1016/j.physa.2007.10.031.

Kästner, K., Caviedes-Voullième, D., Hinz, C. 2025. Formation of spatial vegetation patterns in heterogeneous environments. PLOS One 20, e0324181. doi:10.1371/journal.pone.0324181.

Kästner, K., Van De Vijsel, R.C., Caviedes-Voullième, D., Frechen, N.T., Hinz, C., 2024a. Unravelling the spatial structure of regular dryland vegetation patterns. CATENA 247, 108442. doi:10.1016/j.catena.2024.108442.

Kästner, K., Van De Vijsel, R.C., Caviedes-Voullième, D., Hinz, C., 2024b. A scale-invariant method for quantifying the regularity of environmental spatial patterns. Ecological Complexity 60, 101104. doi:10.1016/j.ecocom.2024.101104.

Kéfi, S., Rietkerk, M., Alados, C.L., Pueyo, Y., Papanastasis, V.P., ElAich, A., De Ruiter, P.C. 2007. Spatial vegetation patterns and imminent desertification in Mediterranean arid ecosystems. Nature 449, 213–217. doi:10.1038/nature06111.

Kéfi, S., Rietkerk, M., Roy, M., Franc, A., De Ruiter, P.C., Pascual, M. 2011. Robust scaling in ecosystems and the meltdown of patch size distributions before extinction: Patch size distributions towards extinction. Ecology Letters 14, 29–35. doi:10.1111/j.1461-0248.2010.01553.x.

Kletter, A.Y., von Hardenberg, J., Meron, E. 2012. Ostwald ripening in dryland vegetation. Communications on Pure & Applied Analysis 11,261–273. doi:10.3934/cpaa.2012.11.261.

Koch, A.J., Meinhardt, H. 1994. Biological pattern formation: from basic mechanisms to complex structures. Reviews of Modern Physics 66, 1481–1507. doi:10.1103/RevModPhys.66.1481.

Krauskopf, B. 2007. Numerical Continuation Methods for Dynamical Systems: Path following and boundary value problems. SpringerLink Bücher,Springer Netherlands, Dordrecht. doi:10.1007/978-1-4020-6356-5.

Lin, Y., Han, G., Zhao, M., Chang, S.X. 2010. Spatial vegetation patterns as early signs of desertification: a case study of a desert steppe in Inner Mongolia, China. Landscape Ecology 25, 1519–1527. doi:10.1007/s10980-010-9520-z.

Maestre, F.T., Escudero, A. 2009. Is the patch size distribution of vegetation a suitable indicator of desertification processes? Ecology 90, 1729–1735. doi:10.1890/08-2096.1.

Maini, P.K., Painter, K.J., Nguyen Phong Chau, H. 1997. Spatial pattern formation in chemical and biological systems. Journal of the Chemical Society, Faraday Transactions 93, 3601–3610. doi:10.1039/a702602a.

Martinez-Garcia, R., Cabal, C., Calabrese, J.M., Hernández-García, E., Tarnita, C.E., López, C., Bonachela, J.A. 2023. Integrating theory and experiments to link local mechanisms and ecosystem-level consequences of vegetation patterns in drylands. Chaos, Solitons & Fractals 166, 112881. doi:10.1016/j.chaos.2022.112881.

McCullen, N., Wagenknecht, T. 2016. Pattern Formation on Networks: from Localised Activity to Turing Patterns. Scientific Reports 6, 27397.doi:10.1038/srep27397.

Meron, E. 2015. Nonlinear Physics of Ecosystems. Taylor & Francis Group. doi:10.1201/b18360.

Meron, E. 2018. From Patterns to Function in Living Systems: Dryland Ecosystems as a Case Study. Annual Review of Condensed Matter Physics 9, 79–103. doi:10.1146/annurev-conmatphys-033117-053959.

Merwin, L., He, T., Lamont, B.B., Enright, N.J., Krauss, S.L. 2012. Low Rate of Between-Population Seed Dispersal Restricts Genetic Connectivity and Metapopulation Dynamics in a Clonal Shrub. PLoS ONE 7, e50974. doi:10.1371/journal.pone.0050974.

Nakao, H., Mikhailov, A.S. 2010. Turing patterns in network-organized activator–inhibitor systems. Nature Physics 6, 544–550. doi:10.1038/nphys1651.

Newman, M.E.J., Watts, D.J. 1999. Scaling and percolation in the small-world network model. Physical Review E 60, 7332–7342. doi:10.1103/PhysRevE.60.7332.

Olesen, J.M., Bascompte, J., Dupont, Y.L., Jordano, P. 2006. The smallest of all worlds: Pollination networks. Journal of Theoretical Biology 240, 270–276. doi:10.1016/j.jtbi.2005.09.014.

Pinto-Ramos, D., Clerc, M.G., Tlidi, M. 2023. Topological defects law for migrating banded vegetation patterns in arid climates. Science Advances 9, eadf6620. doi:10.1126/sciadv.adf6620.

Pinto-Ramos, D., Echeverría-Alar, S., Clerc, M., Tlidi, M. 2022. Vegetation covers phase separation in inhomogeneous environments. Chaos, Solitons & Fractals 163, 112518. doi:10.1016/j.chaos.2022.112518.

Pinto-Ramos, D., Clerc, M.G., Makhoute, A., Tlidi, M. 2025. Aperiodic Clustered and Periodic Hexagonal Vegetation Spot Arrays Explained by Inhomogeneous Environments and Climate Trends in Arid Ecosystems. Geophysical Research Letters 52, e2025GL118462. doi:10.1029/2025GL118462.

Pueyo, Y., Kefi, S., Alados, C.L., Rietkerk, M. 2008. Dispersal strategies and spatial organization of vegetation in arid ecosystems. Oikos 117, 1522–1532. doi:10.1111/j.0030-1299.2008.16735.x.

Rietkerk, M., Bastiaansen, R., Banerjee, S., van de Koppel, J., Baudena, M., Doelman, A. 2021. Evasion of tipping in complex systems through spatial pattern formation. Science 374, eabj0359. doi:10.1126/science.abj0359.

Rietkerk, M., Boerlijst, M., van Langevelde, F., HilleRisLambers, R., De Koppel, J., Kumar, L., Prins, H., De Roos, A. 2002. Self-Organization of Vegetation in Arid Ecosystems. The American Naturalist 160, 524–530. doi:10.1086/342078.

Rietkerk, M., van De Koppel, J. 2008. Regular pattern formation in real ecosystems. Trends in Ecology & Evolution 23, 169–175. doi:10.1016/j.tree.2007.10.013.

Robledo-Arnuncio, J.J., Klein, E.K., Muller-Landau, H.C., Santamaría, L. 2014. Space, time and complexity in plant dispersal ecology. Movement Ecology 2, 16. doi:10.1186/s40462-014-0016-3.

Saixiyala, Chen L., Yi, F., Qiu, X., Sun, H., Cao, H., Baoyin, T., Ye, X., Huang, Z. 2023. Warming in combination with increased precipitation mediate the sexual and clonal reproduction in the desert steppe dominant species Stipa breviflora. BMC Plant Biology 23, 474. doi:10.1186/s12870-023-04439-w.

Scanlon, T.M., Caylor, K.K., Levin, S.A., Rodriguez-Iturbe, I. 2007. Positive feedbacks promote power-law clustering of Kalahari vegetation.Nature 449, 209–212. doi:10.1038/nature06060.

Tamme, R., Götzenberger, L., Zobel, M., Bullock, J.M., Hooftman, D.A.P., Kaasik, A., Pärtel, M. 2014. Predicting species’ maximum dispersal distances from simple plant traits. Ecology 95, 505–513. doi:10.1890/13-1000.1.

Telesford, Q.K., Joyce, K.E., Hayasaka, S., Burdette, J.H., Laurienti, P.J. 2011. The Ubiquity of Small-World Networks. Brain Connectivity 1,367–375. doi:10.1089/brain.2011.0038.

Thomson, F.J., Moles, A.T., Auld, T.D., Kingsford, R.T. 2011. Seed dispersal distance is more strongly correlated with plant height than with seed mass. Journal of Ecology 99, 1299–1307. doi:10.1111/j.1365-2745.2011.01867.x.

Turing, A. 1952. The chemical basis of morphogenesis. Philosophical Transactions of the Royal Society of London. Series B, Biological Sciences 237, 37–72. doi:10.1098/rstb.1952.0012.

Vittoz, P., Engler, R. 2007. Seed dispersal distances: a typology based on dispersal modes and plant traits. Botanica Helvetica 117, 109–124. doi:10.1007/s00035-007-0797-8.

Wang, Z., Xie, L., Prather, C.M., Guo, H., Han, G., Ma, C. 2018. What drives the shift between sexual and clonal reproduction of Caragana stenophylla along a climatic aridity gradient? BMC Plant Biology 18, 91. doi:10.1186/s12870-018-1313-6.

Watts, D.J. 1999. Networks, Dynamics, and the Small-World Phenomenon. American Journal of Sociology 105, 493–527. doi:10.1086/210318.

Watts, D.J., Strogatz, S.H. 1998. Collective dynamics of ‘small-world’ networks. Nature 393, 440–442. doi:10.1038/30918.

Wendel, A.S., Bauke, S.L., Amelung, W., Knief, C. 2022. Root-rhizosphere-soil interactions in biopores. Plant and Soil 475, 253–277. doi:10.1007/s11104-022-05406-4.

Yin, H., Benson, A.R., Leskovec, J. 2018. Higher-order clustering in networks. Physical Review E 97, 052306. doi:10.1103/PhysRevE.97.052306.

Yizhaq, H., Sela, S., Svoray, T., Assouline, S., Bel, G. 2014. Effects of heterogeneous soil-water diffusivity on vegetation pattern formation. Water Resources Research 50, 5743–5758. doi:10.1002/2014WR015362.

Zelnik, Y.R., Meron, E., Bel, G. 2015. Gradual regime shifts in fairy circles. Proceedings of the National Academy of Sciences 112, 12327–12331.doi:10.1073/pnas.1504289112.

Zhang, Y., Zhang, M., Niu, J., Zheng, H. 2016. The preferential flow of soil: A widespread phenomenon in pedological perspectives. Eurasian Soil Science 49, 661–672. doi:10.1134/S1064229316060120.

