## Supplementary Material for "Network-mediated diffusion produces disordered self-organization in vegetation^⋆^"

### S1 Dimensional model

We present here the dimensional version of model (1-2):

$$\frac{\partial B}{\partial t} = \Lambda W B \left(1 - \frac{B}{K}\right)(1 + EB)^2 - MB + D_B \nabla^2 B \tag{S1.1}$$

$$\frac{\partial W}{\partial t} = P - \frac{NW}{1 + \frac{RB}{K}} - \Gamma W B (1 + EB)^2 + D_W \nabla^2 W \tag{S1.2}$$

where $B$ and $W$ refer to biomass and water density (kg/m$^2$), respectively. In equation (S1.1), $\Lambda$ is the rate of biomass growth per water absorbed, $K$ is the maximum standing biomass over a given point, $E$ is the root-to-shoot ratio, $M$ is the plant mortality rate, and $D_B$ is the diffusion constant of biomass. In equation (S1.2), $P$ represents the annual precipitation, $N$ is the evaporation rate, $R$ controls the intensity of evaporation reduction due to shading, $\Gamma$ is the rate of water absorption by plants, and $D_W$ is the water diffusion constant.

Table S1.1 reports the definitions of the dimensionless terms of model (1-2), as well as their values in our simulations.

Table S1.1: Dimensionless terms in equations (1-2), with their relation to the dimensional terms in equations (S1.1-S1.2) and their values in our implementation.

| Dimensionless term | Definition | Value |
|---|---|---|
| $x$ | $X\sqrt{M/D_B}$ | $[0, +\infty)$ |
| $t$ | $MT$ | $[0, +\infty)$ |
| $p$ | $\Lambda P/K\Gamma M$ | $[0, +\infty)$ |
| $b$ | $B/K$ | $[0, 1]$ |
| $w$ | $W\Lambda/K\Gamma$ | $[0, p/\nu]$ |
| $\gamma$ | $K\Gamma/M$ | 0.4571 |
| $\eta$ | $EK$ | 2.8 |
| $\nu$ | $N/M$ | 1.4286 |
| $\rho$ | $R$ | 0.7 |
| $d_w$ | $D_W/D_B$ | 125 |

### S2 Patch coarsening over time

In our model, at intermediate values of $\phi_w$, the coarsening process is incomplete, resulting in stable wide patch size distribution patterns. In figure S2.1 we show the evolution of the average patch size over time for $p = 1.5$, $\phi_w = 0.01$ and $\phi_b = 0$. We see that in the early stages on the simulation the average patch size grows proportionally to $\sqrt{t}$, as observed by Kletter et al. [63].

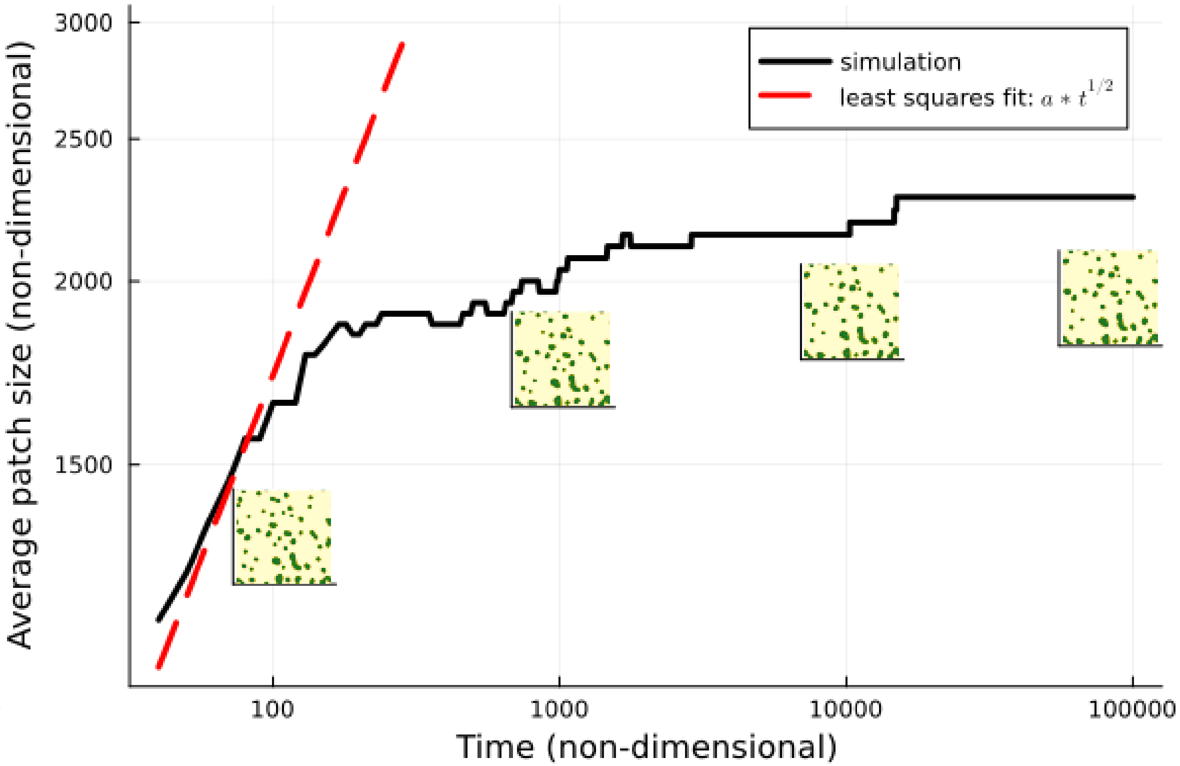


Figure S2.1: Evolution of the average patch size over time on a log-log scale, at $p = 1.5$, $\phi_w = 0.01$ and $\phi_b = 0$. The domain size was 336x336 in non-dimensional units. The red dashed line shows the least mean square fit for a function proportional to $\sqrt{t}$. The insets show snapshots of the patterns reached at different stages of the simulation.

### S3 Effects of changes in isotropic diffusion

In the dimensionless equations (1-2), water and biomass dimensional diffusion constants ($D_W$, $D_B$) appear as their ratio $d_w$. In all the simulations analyzed in the main text, $d_w$ was equal to 125. In order to provide a comparable parameter for measuring the changes in diffusion due to the network structure, we define $\tilde{\phi}$ as the ratio between the number of edges in the biomass and water diffusion networks, so that $\tilde{\phi} = \frac{\phi_w+1}{\phi_b+1}$. We compare the effects of shortcuts in the biomass diffusion network ($\tilde{\phi} < 1$) with those of decreased dimensionless diffusivity ($d_w < 125$), and the effects of shortcuts in the water diffusion network ($\tilde{\phi} > 1$) with those of increased dimensionless diffusivity ($d_w > 125$).

Figure S3.1 shows the results of simulations at 7 different $d_w$ values and 3 different precipitation values. The domain size and simulation time are the same as in figure 3 and 8 of the main text.

Comparing the right side of figure S3.1 ($d_w \geq 125$) with figure 8 of the main text ($\tilde{\phi} > 1$), we notice that the growth in patch size and characteristic wavelength of

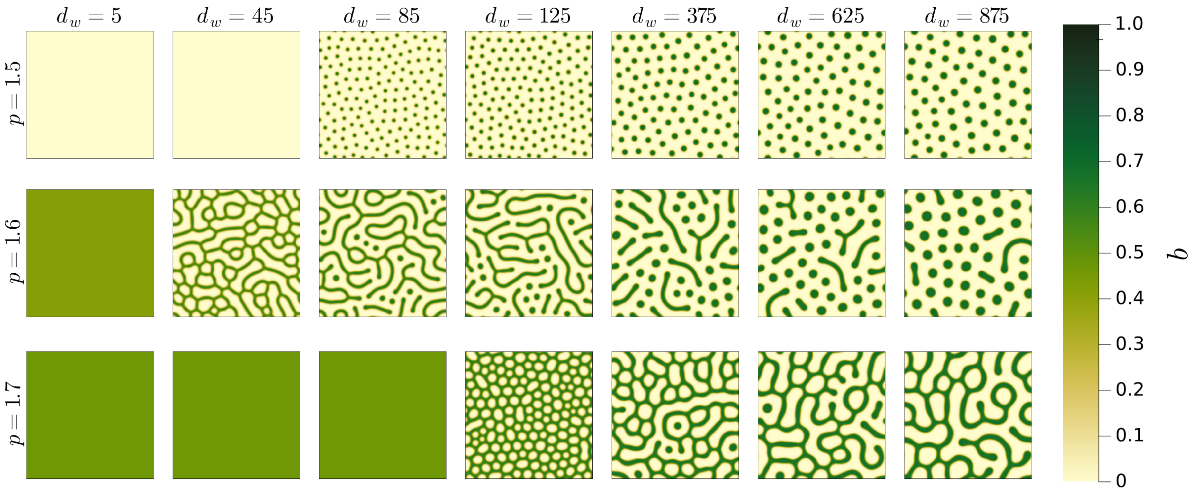


Figure S3.1: Final stage of 21 different simulations at different $d_w$ and $p$ values. The initial condition for all simulations is a uniform state perturbed with uniformly distributed random noise in the interval $[-0.05, 0.05]$. The simulation length is 5000 dimensionless times and the dimensionless domain size is 336x336.

the patterns is common to both an increase in diffusivity and an increase in shortcut density in the water diffusion network. However, increase in diffusivity does not lead to the emergence of irregular and broad patch size patterns as observed in figure 8. Similarly, the low-regularity patterns observed in figure 3 ($\tilde{\phi} < 1$) are not reproduced over a 2-dimensional lattice at low $d_w$ values. The patterns give way to either uniform vegetation or desert, both at low $d_w$ values and at low $\tilde{\phi}$ values.

The relation between the wavelength of the emerging patterns and $d_w$ may also be studied analytically through the dispersion relations of model (1-2), showing the growth rate of different spatial frequencies from a uniform state following a disturbance. Figure S3.2 shows the dispersion relations of the uniform vegetation state at $p = 1.7$, for different $d_w$ values.

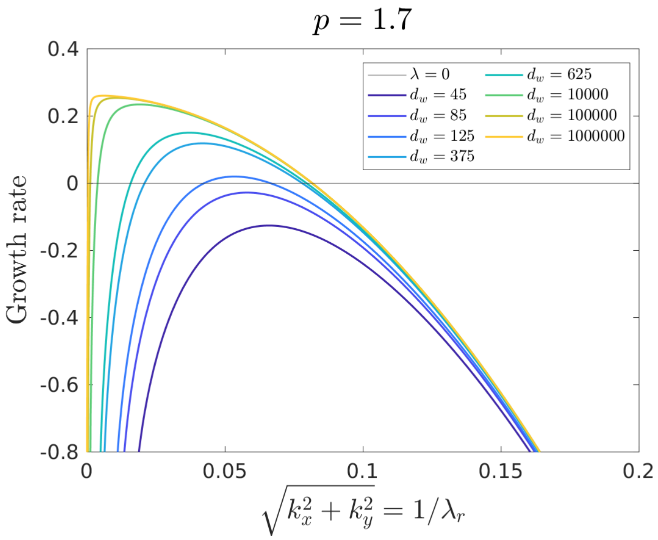


Figure S3.2: Dispersion relations in the local model. We show the growth rate of different spatial frequencies at $p = 1.7$ from a perturbed uniform solution, for different $d_w$ values.

We verify that our simulations are in agreement with the analytical results. All curves are negative at the zero frequency, meaning the uniform solution is marginally stable at $p = 1.7$ (stable to spatially uniform disturbances). The fastest growing spatial frequency decreases for increasing $d_w$ (meaning the characteristic wavelength of the patterns increases), and for $d_w \leq 105$ the uniform vegetation state is stable to all disturbances. We further observe that, at very high diffusivity values, the fastest growing frequency tends towards the lowest non-zero frequency, meaning the non-uniform solution with the longest possible wavelength: a single patch over a bare landscape. Thus, the pattern we observed in the high shortcut density range reflects the theoretical case of $d_w \to \infty$ over a 2-dimensional lattice.

Finally, we consider the behavior of the spatially integrated variables of the patterns in relation to diffusivity over a 2-dimensional lattice, and in relation to shortcut density over an augmented network. Figure S3.3 shows the progression, for varying $d_w$ (with $\tilde{\phi} = 1$), of the average biomass density (figure S3.3a), the vegetation cover fraction (figure S3.3b), the average biomass density in vegetated areas (figure S3.3c), and the average biomass density in bare areas (figure S3.3d). Figure S3.4 shows the same variables for varying $\tilde{\phi}$ (with $d_w = 125$).

For increasing $d_w$, total biomass density and vegetation cover (figures S3.3a and S3.3b) increase at low precipitation values and decreases high precipitation values, resulting in a lower sensitivity of the patterns to precipitation, similarly to the behavior over an augmented network, as shown in figures S3.4a and S3.4b. Conversely, for $d_w \leq 125$, the patterned states separate into colonization and retreat patterns, as discussed in section 3.1 of the main text, growing more distant from each other in terms of average biomass density and vegetation cover, up to the point where the system falls into either desert (for $p < 1.52$) or uniform vegetation state (for $p \geq 1.52$). This is again qualitatively similar to what is shown on the left side of figures S3.4a and S3.4b.

Higher values of $d_w$ lead to patterns with a higher "contrast" between vegetated and bare regions, as the biomass density increases in the vegetated areas (figure S3.3c and S3.3d). For $d_w \leq 125$, the patterns tend towards spatial homogenization, either through a "colonization" process (biomass increases in the bare areas for decreasing $d_w$) or a "retreat" process (biomass decreases in the bare areas for decreasing $d_w$) as shown in figure S3.3d. In figure S3.4c and S3.4d we observe a similar behavior in relation to $\tilde{\phi}$, as previously discussed in section 3.1 and 3.2.

Hence, we may deduce that the qualitative behavior of the mean-field variables in relation to the shortcut densities may be explained as a response to the facilitated flow of biomass/water across the system. However, as evidenced by the steep gradients in figure S3.4, and by the single patch patterns reached at $\phi_w \sim 0.1$ (see main text, figure 8), the system appears much more sensitive to the addition of shortcuts than to the increase in isotropic diffusion over a 2-dimensional lattice. This is reasonable if we consider that the addition of a random link increases flow not only through the random link itself but across the whole system as a result of the increased connectivity. The average shortest path length, defined as the minimum number of edges to be crossed in

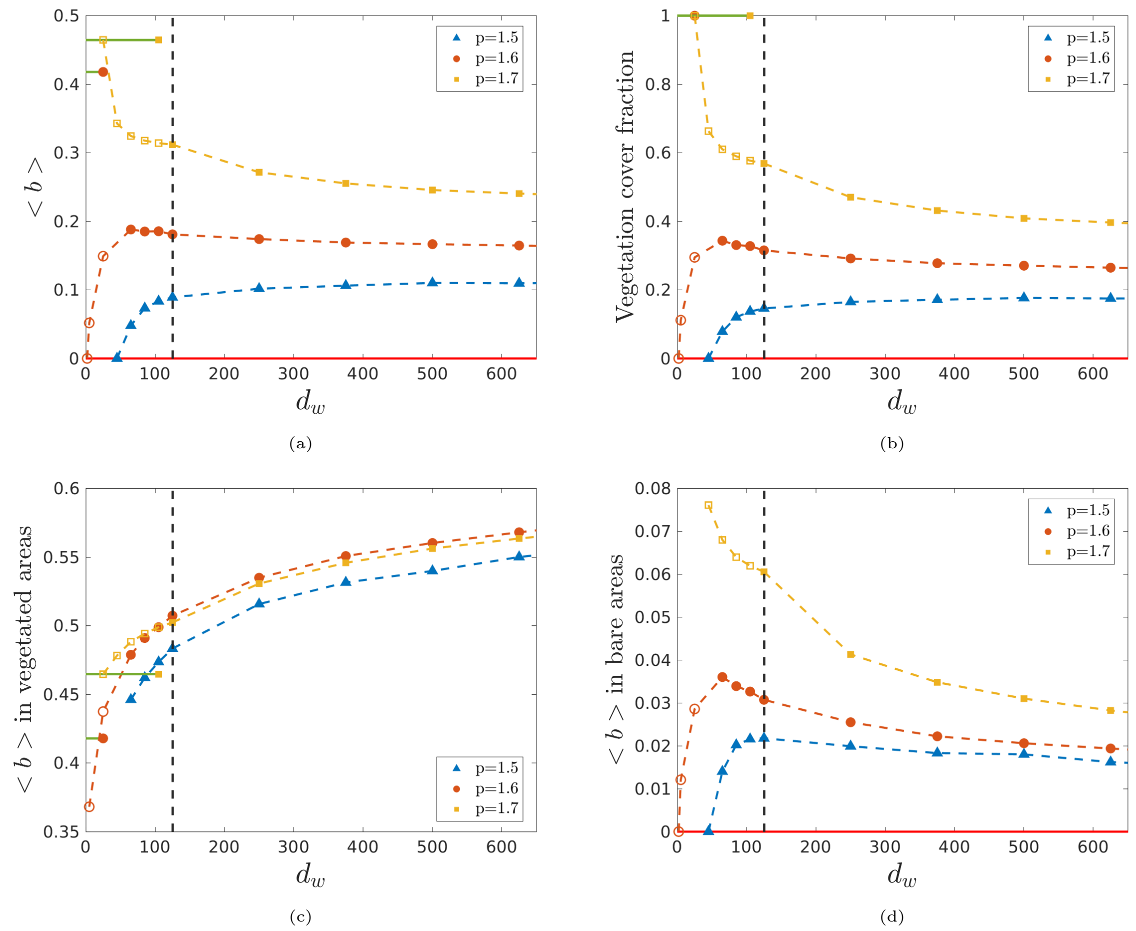

Figure S3.3: Behavior of the mean-field variables in relation to $d_w$. Average biomass density over the whole domain (S3.3a), fraction of the domain where $b > 0.1$ (S3.3b), average biomass density in vegetation-covered areas ($b > 0.1$) (S3.3c), average biomass density in bare areas ($b \leq 0.1$) (S3.3d), . The dashed black line indicates $d_w = 125$. Asterisks refer to uniform states.

order to connect two nodes of the network, has a strongly non-linear relation with the number of shortcuts, as discussed in section 4.2.

### S4 Full derivation of equations (6-7) (mean-field approximations)

Starting from equations (3-4):

$$\frac{\partial b_i}{\partial t} = \gamma w_i b_i (1 - b_i)(1 + \eta b_i)^2 - b_i + \frac{1}{\Delta x^2}\left(\sum_{j=1}^{N} A^b_{i,j} b_j - k^b_i b_i\right) \qquad \text{(S4.1)}$$

$$\frac{\partial w_i}{\partial t} = p - \frac{\nu w_i}{1 + \rho b_i} - \gamma w_i b_i (1 + \eta b_i)^2 + \frac{d_w}{\Delta x^2}\left(\sum_{j=1}^{N} A^w_{i,j} w_j - k^w_i w_i\right) \qquad \text{(S4.1)}$$

We may rewrite the adjacency matrices $\mathbf{A^b}$ and $\mathbf{A^w}$ and the degree vectors $\mathbf{k^b}$ and $\mathbf{k^w}$ as the sum of a regular lattice component ($\hat{\mathbf{A}}^\mathbf{b}, \hat{\mathbf{A}}^\mathbf{w}, \hat{\mathbf{k}}^\mathbf{b}, \hat{\mathbf{k}}^\mathbf{w}$) and a shortcut component ($\tilde{\mathbf{A}}^\mathbf{b}, \tilde{\mathbf{A}}^\mathbf{w}, \tilde{\mathbf{k}}^\mathbf{b}, \tilde{\mathbf{k}}^\mathbf{w}$). So, for each pair of nodes $i, j$:

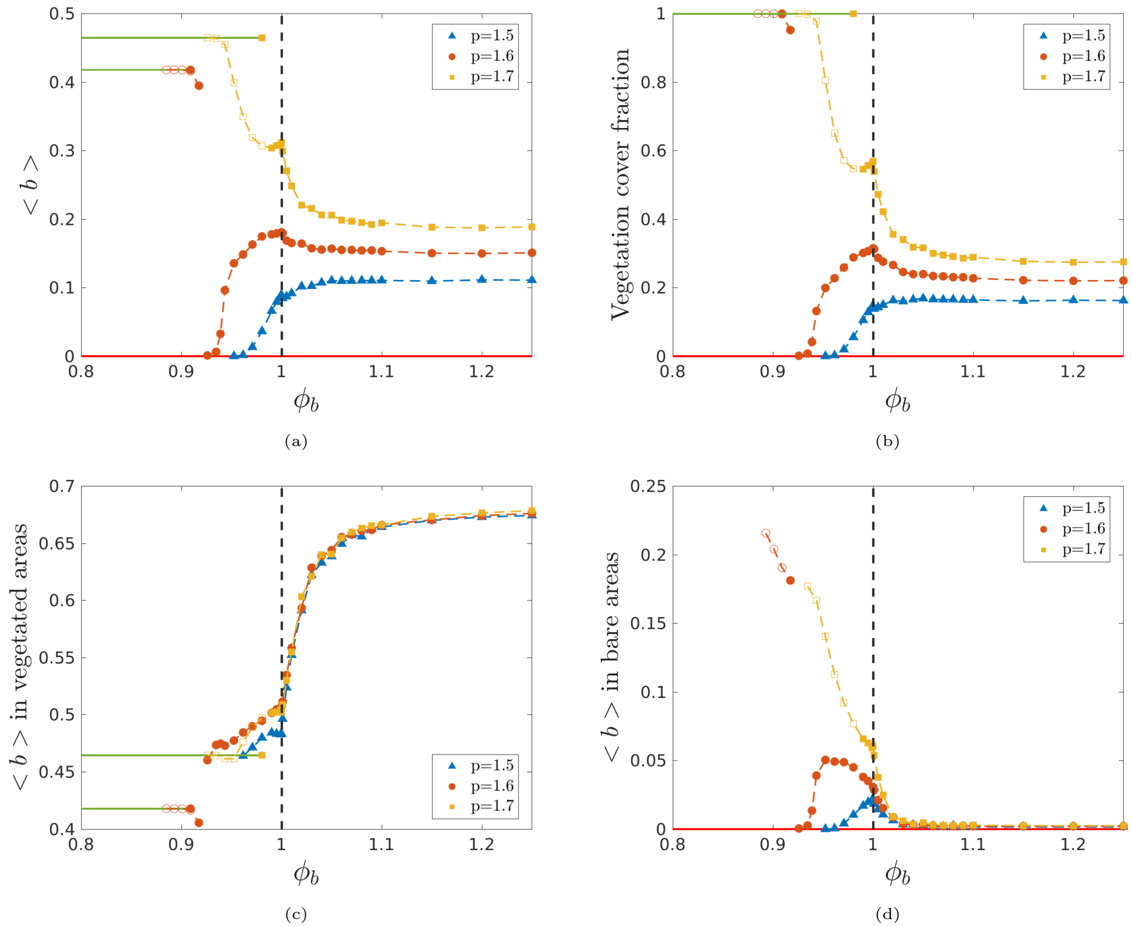


Figure S3.4: Behavior of the mean-field variables in relation to $\tilde{\phi}$. Average biomass density over the whole domain (S3.4a), fraction of the domain where $b > 0.1$ (S3.4b), average biomass density in vegetation-covered areas ($b > 0.1$) (S3.4c), average biomass density in bare areas ($b \leq 0.1$) (S3.4d). The dashed black line indicates $\tilde{\phi} = 1$ ($\phi_w = \phi_b = 0$). Asterisks refer to uniform states.

$$A^b_{i,j} = \hat{A}^b_{i,j} + \tilde{A}^b_{i,j}(\phi_b) \tag{S4.3}$$

$$A^w_{i,j} = \hat{A}^w_{i,j} + \tilde{A}^w_{i,j}(\phi_w) \tag{S4.4}$$

$$k^b_i = \hat{k}^b_i + \tilde{k}^b_i(\phi_b) \tag{S4.5}$$

$$k^w_i = \hat{k}^w_i + \tilde{k}^w_i(\phi_w) \tag{S4.6}$$

Inserting into equations (S4.1-S4.2), one obtains

$$\frac{\partial b_i}{\partial t} = \gamma w_i b_i (1 - b_i)(1 + \eta b_i)^2 - b_i + \frac{1}{\Delta x^2}\left(\sum_{j=1}^{N} \hat{A}^b_{i,j} b_j - \hat{k}^b_i b_i + \sum_{j=1}^{N} \tilde{A}^b_{i,j} b_j - \tilde{k}^b_i b_i\right) \tag{S4.7}$$

$$\frac{\partial w_i}{\partial t} = p - \frac{\nu w_i}{1+\rho b_i} - \gamma w_i b_i (1 + \eta b_i)^2 + \frac{d_w}{\Delta x^2}\left(\sum_{j=1}^{N} \hat{A}^w_{i,j} w_j - \hat{k}^w_i w_i + \sum_{j=1}^{N} \tilde{A}^w_{i,j} w_j - \tilde{k}^w_i w_i\right) \tag{S4.8}$$

Preserving diffusion over the regular lattice, we approximate the biomass and water density values of each node $j$ connected to $i$ through a shortcut to the domain averages ($\langle b\rangle$ and $\langle w\rangle$). Considering $\sum_{j=1}^{N}\tilde{A}^{w}_{i,j}=\tilde{k}^{w}_{i}$ and $\sum_{j=1}^{N}\tilde{A}^{b}_{i,j}=\tilde{k}^{b}_{i}$, we have:

$$\frac{\partial b_i}{\partial t}=\gamma w_i b_i(1-b_i)(1+\eta b_i)^2-b_i+\frac{1}{\Delta x^2}\left(\sum_{j=1}^{N}\hat{A}^{b}_{i,j}b_j-\hat{k}^{b}_{i}b_i+\tilde{k}^{b}_{i}(\langle b\rangle-b_i)\right) \tag{S4.9}$$

$$\frac{\partial w_i}{\partial t}=p-\frac{\nu w_i}{1+\rho b_i}-\gamma w_i b_i(1+\eta b_i)^2+\frac{d_w}{\Delta x^2}\left(\sum_{j=1}^{N}\hat{A}^{w}_{i,j}w_j-\hat{k}^{w}_{i}w_i+\tilde{k}^{w}_{i}(\langle w\rangle-w_i)\right) \tag{S4.10}$$

Secondly, we approximate the number of points connected to $i$ through shortcuts ($\tilde{k}^{b}_{i}(\phi_b)$, $\tilde{k}^{w}_{i}(\phi_w)$) to the average value over all nodes $i$ of the network. As noted in section 2.3, in our network model, $\phi$ stands for the ratio between the number of added shortcuts and the number of links of the regular lattice. Thus, over a lattice of $N$ nodes with periodic boundary conditions, we add $2\phi N$ shortcuts. Since each shortcut adds an extra neighbor to two nodes, the average number of extra neighbors per node is $4\phi$. We apply $\tilde{k}^{b}_{i}(\phi_b)\approx 4\phi_b$ and $\tilde{k}^{b}_{i}(\phi_w)\approx 4\phi_w$ and obtain equations:

$$\frac{\partial b_i}{\partial t}=\gamma w_i b_i(1-b_i)(1+\eta b_i)^2-b_i+\frac{1}{\Delta x^2}\left(\sum_{j=1}^{N}\hat{A}^{b}_{i,j}b_j-\hat{k}^{b}_{i}b_i+4\phi^{mf}_{b}(\langle b\rangle-b_i)\right) \tag{S4.11}$$

$$\tfrac{\partial w_i}{\partial t}=p-\tfrac{\nu w_i}{1+\rho b_i}-\gamma w_i b_i(1+\eta b_i)^2+\tfrac{d_w}{\Delta x^2}\left(\textstyle\sum_{j=1}^{N}\hat{A}^{w}_{i,j}w_j-\hat{k}^{w}_{i}w_i+4\phi^{mf}_{w}(\langle w\rangle-w_i)\right) \tag{S4.12}$$

in which we have redefined $\phi_b$ and $\phi_w$ and $\phi^{mf}_{b}$ and $\phi^{mf}_{w}$ to distinguish between the two models.

### S5 Additional figures for the analysis of the mean-field model

Figure S5.1 shows the comparison between the patterns produced through the mean-field approximation and the ones produced through the full model, in terms of their spatially integrated variables. As described in the main text, the approximation appears valid in the high and the low shortcut density limits ($\phi_w \gtrsim 0.1$ and $\phi_w \lesssim 0.001$), but fails in the intermediate range.

Figure S5.2 shows the growth rate of different spatial frequencies from a perturbed uniform vegetation solution in the mean-field model. We observe a behavior consistent with our numerical simulations. For increasing $\phi^{mf}_{b}$, the fastest growing frequency remains constant but maximum growth rate decreases, and for $\phi^{mf}_{b} > 0.005$ the uniform solution is stable. Conversely, for increasing $\phi^{mf}_{w}$ the fastest growing frequency decreases and for $\phi^{mf}_{w} > 0.01$ the lowest available frequency (single patch state) has non-zero growth rate, leading to the slow coarsening process.

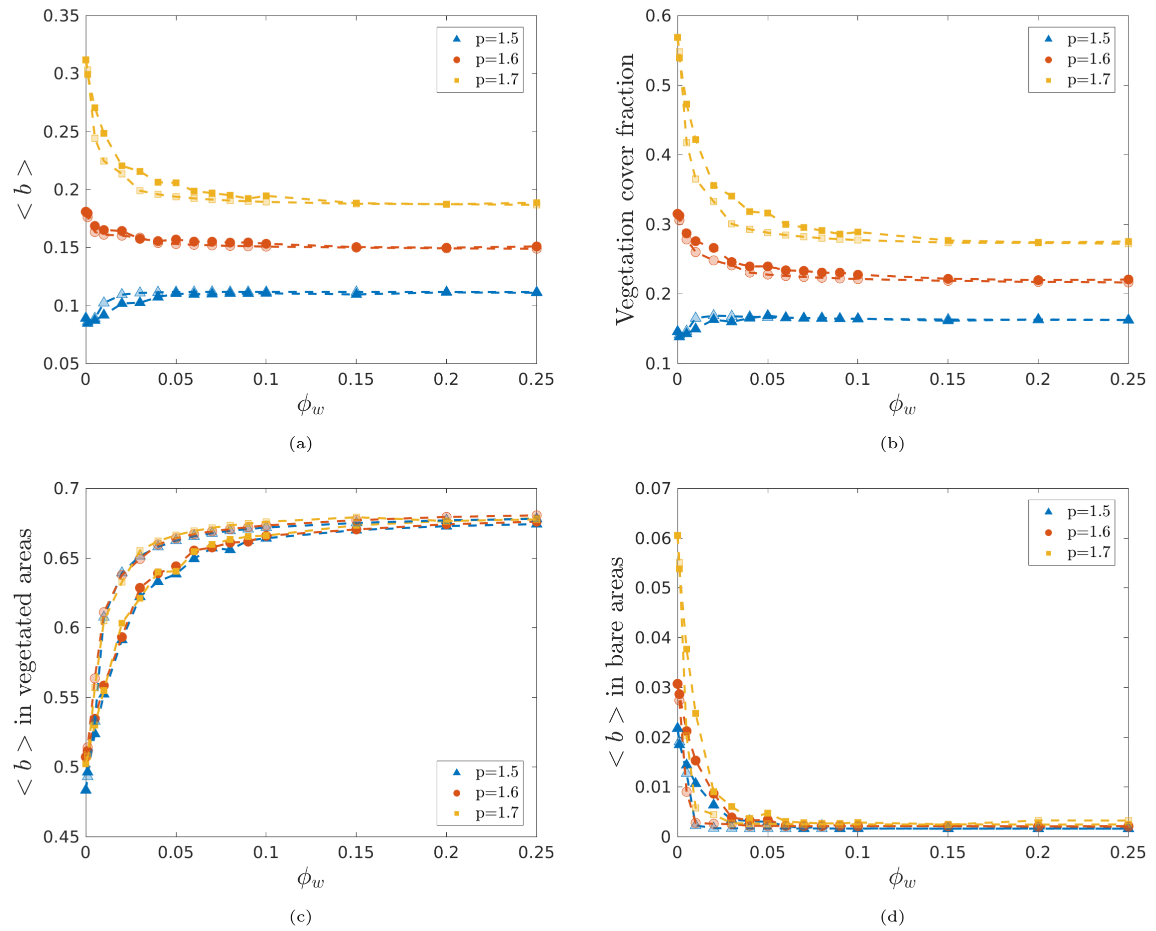


Figure S5.1: Behavior of the spatially integrated variables in relation to $\phi_w$ on the full model (filled scatter points points), and in relation to $\phi_w^{mf}$ in mean-field approximation (non-filled scatter points). The different plots show: average biomass density over the whole domain (a), fraction of the domain where $b > 0.22$ (b), average biomass density in vegetation-covered areas ($b > 0.22$) (c), average biomass density in bare areas ($b \leq 0.22$) (d).

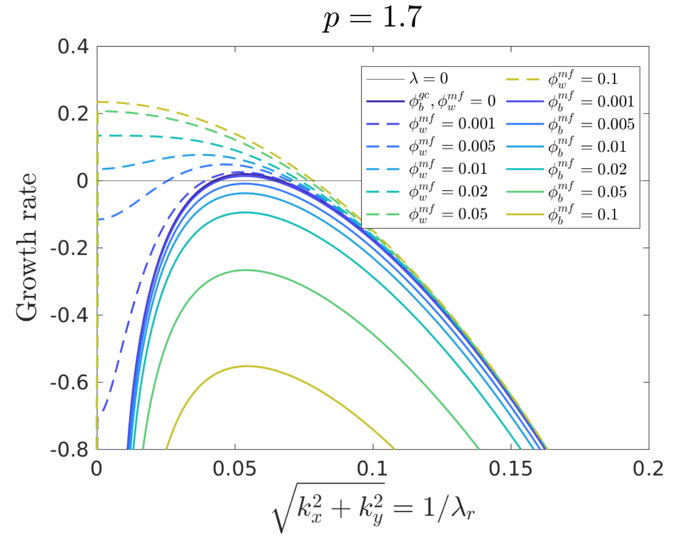


Figure S5.2: Dispersion relations in the mean-field model. We show the growth rate of different spatial frequencies at $p = 1.7$ from a perturbed uniform vegetation solution, for different $\phi_w^{mf}$ and $\phi_b^{mf}$ values.
